# A Functional Map of the Human Intrinsically Disordered Proteome

**DOI:** 10.1101/2024.03.15.585291

**Authors:** Iva Pritišanac, T. Reid Alderson, Đesika Kolarić, Taraneh Zarin, Shuting Xie, Alex Lu, Aqsa Alam, Abdullah Maqsood, Ji-Young Youn, Julie D. Forman-Kay, Alan M. Moses

## Abstract

Intrinsically disordered regions (IDRs) represent at least one-third of the human proteome and defy the established structure-function paradigm. Because IDRs often have limited positional sequence conservation, the functional classification of IDRs using standard bioinformatics is generally not possible. Here, we show that evolutionarily conserved molecular features of the intrinsically disordered human proteome (IDR-ome), termed evolutionary signatures, enable classification and prediction of IDR functions. Hierarchical clustering of the human IDR-ome based on evolutionary signatures reveals strong enrichments for frequently studied functions of IDRs in transcription and RNA processing, as well as diverse, rarely studied functions, ranging from sub-cellular localization and biomolecular condensates to cellular signaling, transmembrane transport, and the constitution of the cytoskeleton. We exploit the information that is encoded within evolutionary conservation of molecular features to propose functional annotations for every IDR in the human proteome, inspect the conserved molecular features that correlate with different functions, and discover frequently co-occurring IDR functions on the proteome scale. Further, we identify patterns of evolutionary conserved molecular features of IDRs within proteins of unknown function and disease-risk genes for conditions such as cancer and developmental disorders. Our map of the human IDR-ome should be a valuable resource that aids in the discovery of new IDR biology.

## Introduction

The sequence-structure-function paradigm in molecular biology posits that the amino-acid sequence of a protein encodes its three-dimensional structure, which determines the function of the protein. The close relationships between sequence, structure, and function are routinely exploited to infer function from sequence or structural data (Ashburner et al. 2000; Lee et al. 2007; Radivojac et al. 2013; Sanderson et al. 2023; Yu et al. 2023b), trace the evolutionary history of protein-protein interactions (Steube et al. 2023), design *de novo* proteins with desired folds or functions (Huang et al. 2016; Kuhlman & Bradley 2019; Yeh et al. 2023), and predict the pathogenicity of sequence variants in the human genome (Adzhubei et al. 2010; Frazer et al. 2021; Hopf et al. 2017; Luppino et al. 2023). Indeed, structural information recovered from amino-acid sequence alignments is central to state-of-the-art protein structure prediction methods (Baek et al. 2021; Jumper et al. 2021). However, the sequence-structure-function paradigm does not apply to the approximately one-third of residues in the human proteome that map to intrinsically disordered regions (IDRs), which lack stable secondary and tertiary structure and exhibit poor positional sequence conservation (Forman-Kay & Mittag 2013; Van Der Lee et al. 2014; Wright & Dyson 2015). Despite their lack of ordered structural elements, IDRs function in key cellular processes (Holehouse & Kragelund 2023) and frequently act as hubs in protein-protein interaction networks (Tompa et al. 2014), often via transient, multivalent interactions that promote phase separation and involvement in biomolecular condensates (Borcherds et al. 2021).

While the presence of IDRs in proteins can generally be predicted with high accuracy from their amino-acid sequences (Emenecker et al. 2021; Necci et al. 2021), identifying the relationship between the sequences and functions of IDRs remains a difficult task (Basu et al. 2023; Chow et al. 2023; Hu et al. 2021; Lu et al. 2022; Pang & Liu 2022; Pritišanac et al. 2019; Zarin et al. 2019, 2021; Zhao et al. 2021). Focusing on the segments of IDR sequences that show strong similarity in sequence alignments (which we refer to as “positional conservation”) has provided rich insights into the functions of so-called short-linear motifs (SLiMs) and Molecular Recognition Features (MoRFs) (Davey et al. 2023; Kumar et al. 2022; Malhis & Gsponer 2015; Mohan et al. 2006; Tompa et al. 2014). However, positionally conserved elements typically constitute only a minor fraction of an IDR sequence, and many of the experimentally characterized SLiMs are not positionally conserved (Davey et al. 2012; Kumar et al. 2022; Nguyen Ba et al. 2012; Van Roey et al. 2014). Although we and others showed that approximately 15% of human IDRs contain significant positional alignment due to the acquisition of a conditional fold in particular functional contexts (Alderson et al. 2023; Piovesan et al. 2022), it is appreciated that the majority of positions in the sequences of IDRs appear to evolve more rapidly relative to ordered regions in the same proteins (Brown et al. 2002; Davey et al. 2012). Rapid evolution in IDRs reflects the absence of stable folded structure, since positional conservation is directly linked to evolutionary pressure to maintain a three-dimensional fold (Pritišanac et al. 2019). Thus, because IDRs exhibit limited positional conservation in multiple sequence alignments, standard bioinformatic approaches that rely on these alignments can only provide limited insight into the functional classification of IDRs (Pritišanac et al. 2019; Zarin et al. 2019, 2021). For intrinsically disordered proteins (IDPs), which are fully disordered and make up ∼5% of the human proteome (*ca.* 1000 proteins) (Tsang et al. 2020), predictions of function are even more limited due to the lack of any folded domains (Basu et al. 2023).

The functional importance of IDRs and IDPs is increasingly appreciated, especially in the context of phase separation (Alberti & Dormann 2019; Basu et al. 2020; Mensah et al. 2023; Molliex et al. 2015; Nakamura et al. 2023; Patel et al. 2015). Furthermore, IDRs are often dysregulated in diseases such as cancer, amyotrophic lateral sclerosis, and other neurological disorders (Alberti & Dormann 2019; Tsang et al. 2020; Uversky et al. 2008), with increasing reports of disease-associated sequence variants that map to IDRs (Alderson et al. 2021; Mensah et al. 2023; Vacic et al. 2012). The interpretation of effects of mutations in IDRs on protein function is limited, as most utilized variant effect predictors compute effects on fold stability and other structural features, e.g., changes to enzyme active sites or interfaces (Backwell & Marsh 2022). Thus, an understanding of how the sequences of human IDRs relate to biological function is urgently needed.

The overall importance of IDRs in health and disease has stimulated efforts to predict their biological function without relying on multiple sequence alignments (Cohan et al. 2022; Lancaster et al. 2014; Langstein-Skora et al. 2022; Pang et al. 2024; Shinn et al. 2022; Staller et al. 2018, 2022; Vernon et al. 2018; Zarin et al. 2017, 2019, 2021). IDRs generally show strong evolutionary conservation of sequence-derived molecular features that are not positionally constrained (Alston et al. 2023; Beh et al. 2012; González-Foutel et al. 2022; Staller et al. 2022; Zarin et al. 2017, 2019, 2021). In a series of recent studies, we showed how evolutionary properties of bulk molecular features that are computable from IDR sequences can be used to cluster and classify yeast IDRs into an unexpectedly large number of functional groups (Zarin et al. 2019, 2021).

Here, we show that human IDRs are amenable to systematic functional classification based on a broad set of bulk molecular features that are readily computable from IDR sequences. We provide, to our knowledge, the first comprehensive functional map of IDRs within the human proteome (IDR-ome). We obtain estimates for the proportion of human IDRs associated with various functions, and train classifiers to predict functions for unannotated IDPs and IDRs from sequence alone. Since the functional map of human IDRs is based on evolutionary conservation of simple molecular features, we can determine which features are associated with different groups of IDRs, such as those involved in the formation of biomolecular condensates or those associated with disease-risk genes. We expect that the patterns of conservation of molecular features, together with the functional map of IDRs, will represent a critical resource for generating testable hypotheses for experimental research for biochemists and cell biologists.

## Results

### Intrinsic disorder is abundant in human proteins

To build a global functional map of the human IDRs, we first identified the boundaries of IDRs in the human proteome using a state-of-the-art bioinformatics tool, SPOT-Disorder (SPOTD) (Hanson et al. 2017) (**Methods**). SPOTD predicts that *ca.* 32.8% of the residues in the human proteome are disordered, similar to other reports (Necci et al. 2021). We then filtered the predictions to keep only regions of 30 or more consecutive disordered residues (**Methods**), henceforth referred to as intrinsically disordered regions or IDRs and comprising a total of 21,252 sequences (Tsang et al. 2020). Nearly 60% of proteins in the human proteome contain at least one IDR (**Figure 1A**). Predominantly disordered proteins, or those that contain more than 50% disordered residues, amount to nearly 20% of the proteome, whereas entirely disordered proteins (*i.e.*, IDPs) account for 5% (**Figure 1A**). The IDR lengths follow a power-law distribution with a median IDR length of 74 residues (**Figure 1B**). Approximately 90% of the human IDRs fall between 30 and 200 residues in length. However, the human proteome also contains some very long IDRs, with 280 IDRs or IDPs having more than 1000 consecutively disordered residues (**Figure 1B**).

**Figure 1.**
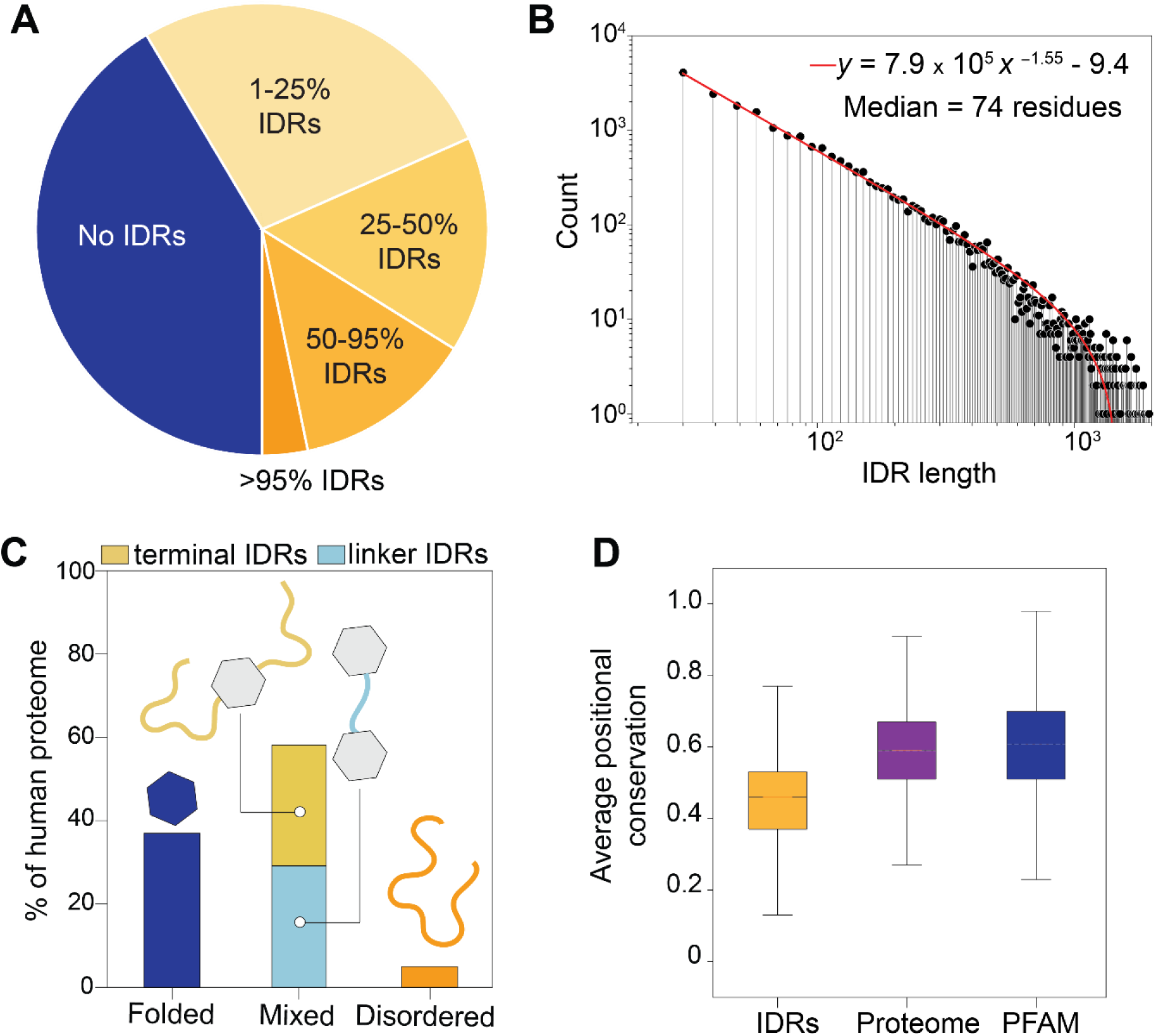
Extent and distribution of intrinsic disorder in the human proteome. (**A)** Break-up of the human proteome by the protein intrinsic disorder content based on the SPOT-Disorder predictor (Hanson J et al. 2017 & 2018). A minimum of 30 consecutive disordered residues was used as a criterion to define an IDR. The pie chart is stratified to emphasize the extent of intrinsically disordered regions as a percentage of the total protein sequence, i.e., 50-95% means that between 50% and 95% of the entire protein sequence is intrinsically disordered. It is evident that the majority of proteins in the human proteome have IDRs, with a sizeable fraction (15%) containing more than half of the sequence in disordered regions. (**B**) Distribution of lengths of human IDRs. The IDR lengths follow a power-law distribution (*A*x*^b^* + *c*), with *A*, *b*, and *c* as given on the plot. (**C**) Distribution of intrinsic disorder in the human proteome. Fully disordered and folded proteins are shown in green and pink, respectively. Mixed proteins that contain both IDRs and folded domains are separated into terminal IDRs (yellow) that are located at the N-or C-termini of a protein, and linker IDRs (blue) that are between folded domains. (**D**) Comparison of sequence similarity in alignments as measured by the average Jensen-Shannon divergence over alignment columns with the background distribution given as observed in extant human IDR sequences. The average positional conservation when aligned to ENSEMBL orthologs is shown in a box plot for IDRs, the entire human proteome, and PFAM domains (Mistry J, *et al*. 2021). Note that gaps, which are more frequent in IDRs when compared to PFAM domains (Khan T, *et al*. 2015, Chow CFW, *et al*. 2023), were ignored in the computation of positional conservation and thus the conservation for IDRs as shown is an overestimate. The boxes extend to the upper and lower quartiles, and the lines within the boxes corresponds to the median value.

We also checked if IDRs are more likely to be located on the N-or C-terminal regions of a protein (termini) or between folded domains (linkers). Approximately equal percentages of terminal and linker IDRs were identified (**Figure 1C**). Thus, most IDRs in the human proteome are of medium length (30-200 residues) and exist alongside proteins that also harbor folded domains. We also confirmed that, as expected, the predicted IDRs show generally lower levels of positional sequence-similarity in alignments of homologs from the Ensembl database (Howe et al. 2021) as compared to folded protein domains (**Figure 1D**). Further, most human IDRs are not easily assigned to protein families using sequence alignments, and only 21% of human IDRs show significant sequence similarity (BLAST E-value < 1e-6) to any another IDR in the human proteome (**Supplementary** Figure 1). Of the IDRs that do show sequence similarity, over 80% (50%) of these IDRs have five or fewer (one) BLAST hits (**Supplementary** Figure 1), suggesting that the sequence homology of any IDR is restricted to a small number of related IDRs, usually members of a small gene family. Interestingly, we observe that 44% of fully disordered proteins (IDPs) have BLAST hits to any other IDR in the proteome (**Supplementary** Figure 1), which suggests that IDPs are more likely to fall into small gene families compared to IDRs.

### A global map of the human IDR-ome based on evolutionary conservation of molecular features

Following previous work (Zarin et al. 2019) we next sought to summarize IDRs using conservation of bulk molecular features (**Methods**, **Supplementary Table 1**). We first compiled a comprehensive list of 147 bulk molecular features that have been shown to be important for function of IDRs in different studies, including known short linear interaction motifs (SLiMs), physicochemical properties (e.g. hydrophobicity, polarity, charge, charge patterning), residue composition, and (homo-)repeats (Chavali et al. 2017, 2020; Gemayel et al. 2015; Kumar et al. 2022; Mao et al. 2010; Ravarani et al. 2018; Schlessinger et al. 2011; Strickfaden et al. 2007; Warren & Shechter 2017).

We next developed a computational protocol, conceptually based on (Zarin et al. 2019), to estimate evolutionary conservation of molecular features in the human IDR-ome without relying on conventional multiple sequence alignments (**Supplementary** Figure 2). Briefly, we assessed the evolutionary conservation of molecular features in the human IDR-ome by comparing observed sets of homologs to simulations of evolution of IDRs (Zarin et al. 2019). We made technical changes to a previously applied approach (Zarin et al. 2019) Zarin et al. 2019, 2021) to increase the computational efficiency so that we could apply it to the larger and more complex human IDR-ome (see **Methods**, **Supplementary** Figure 3, **Supplementary** Figure 4). We use standard Z-scores to compare the observed distributions of molecular features in homologous IDR sequences to those expected from simulations of IDR sequences under a null hypothesis, which assumes no evolutionary conservation of molecular features (**Figure 2**, **Supplementary** Figure 2). We refer to a set of Z-scores for all molecular features as an evolutionary signature of an IDR, which represents the pattern of conserved molecular features. Positive Z-scores indicate a feature value greater than expected from the null hypothesis (the simulations), while negative Z-scores indicate a feature value smaller than expected (**Supplementary** Figure 2). Negative Z-scores can suggest either depletion of a feature (e.g., selection against hydrophobic residues) or a strongly negative value (e.g., selection for a net charge far below the expectation).

**Figure 2.**
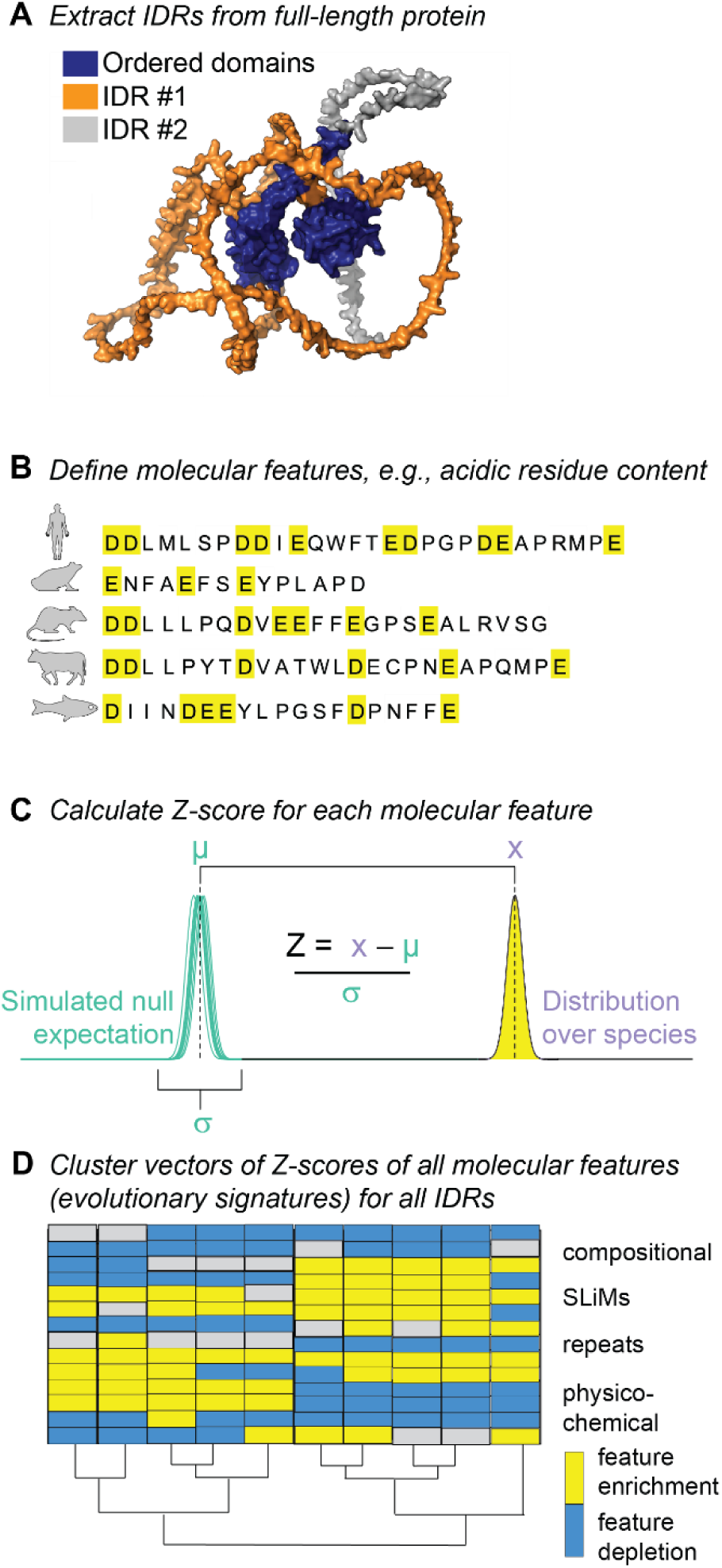
Computing evolutionary signatures in human IDRs. (**A**) A sequence-based disorder predictor is used to define boundaries of IDRs more than 30 consecutive residues in length. The AlphaFold2 structural model of p53 is shown here to illustrate the IDRs (orange, grey) and folded domains (blue). (**B**) Over 140 molecular features are computed for each human IDR sequence and for IDR sequences from other species in the set of orthologues. In the illustrated example, the content of acidic residues is the molecular feature that is computed. (**C**) A Z-score is computed between the evolutionary mean of feature in extant species (*x*) and the average evolutionary mean of the feature in simulations of the null-hypothesis (μ), normalized by the standard deviation of the evolutionary mean in simulations of the null-hypothesis (σ). (**D**) Cluster vectors of Z-scores of all molecular features (evolutionary signatures) for all human IDRs. The resultant evolutionary signatures summarize conservation across the IDRome. When IDRs are hierarchically clustered based on these Z-scores, patterns emerge that define a global map of the IDRome in which IDRs with similar evolutionary signatures appear in close proximity.

We computed evolutionary signatures for 19,032 human IDRs (see Methods) and clustered the IDR-ome (see **Methods**) to identify groups of IDRs that share patterns of conservation (**Figure 3A**, **Supplementary** Figure 5). In this global map of the IDR-ome (**Figure 3A**), IDRs that have similar evolutionary signatures are placed closer to one another. We hypothesize that similarity in the evolutionary patterns of molecular features is analogous to sequence similarity detected in alignments for folded protein regions (e.g., by using PSI-BLAST (Altschul et al. 1997)). Our map reveals a large number of clusters of IDRs, which are defined by distinct patterns of conserved molecular features, i.e., evolutionary signatures (**Figure 3**, **Supplementary** Figure 5).

**Figure 3.**
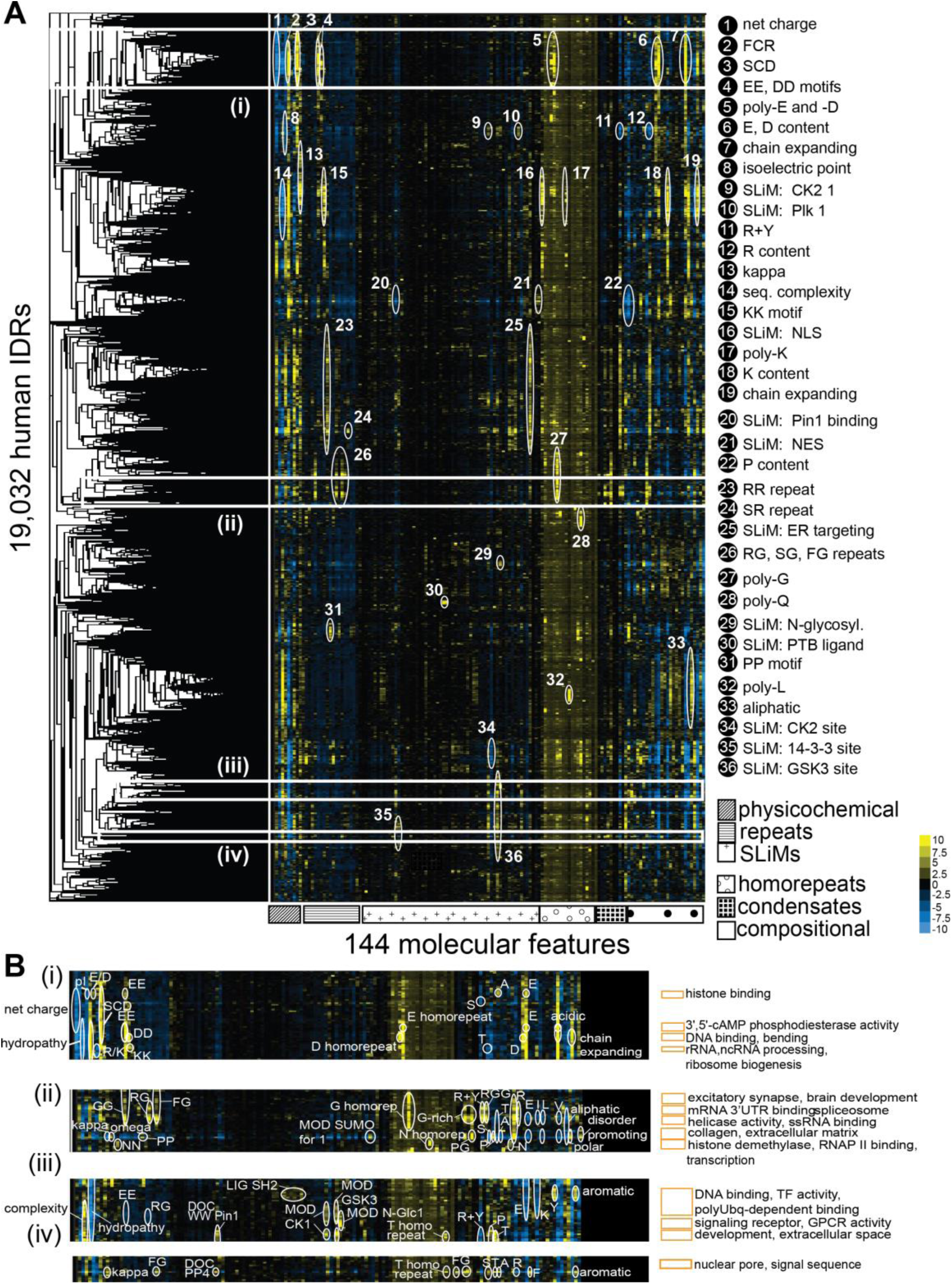
A global map of human IDRs obtained through clustering of evolutionary signatures. **(A)** Hierarchical clustering of 19,032 human IDRs (y-axis) based on the evolutionary conservation of 144 different molecular features (x-axis). The molecular features are grouped into six different categories (physicochemical, repeats, SLiMs, homorepeats, condensates, or compositional biases). This global map of the human IDR-ome shows conservation Z-scores, with some of the dominant molecular features annotated with white circles and numbers, described in the legend (right). A positive or negative Z-score, respectively, is defined by a higher or lower value of a mean of a molecular feature over orthologous IDRs than expected based on a simulation of an absence of evolutionary conservation. White rectangles indicate areas of selected clusters featured in panel B. **(B)** Clusters that are defined by strong patterns of Z-scores often contain a statistically significant overrepresentation of GO-term molecular functions, biological processes and/or sub cellular localizations, as listed here for select examples in areas **(i)**, **(ii)**, **(iii)** and **(iv)** from the panel A. A detailed view of statistically overrepresented terms and features for the rest of the map are available in **Supplementary Figure S5**. Complete information on statistics of functional overrepresentation associated with each cluster selected manually or extracted using an automatic protocol are available in **Supplementary Table 2.**

### Assigning biological functions to clusters of IDRs

Our global map of the IDR-ome is based on evolutionary conserved molecular features that could be important for specific functions of IDRs. To test if the patterns of conservation of molecular features are associated with specific functions, we performed overrepresentation analysis for the Gene Ontology (GO) annotations of the proteins in which IDRs were found (**Methods**). We manually selected 93 clusters from the map, focusing on patterns of Z-score signals. Among these clusters, 53 (i.e., 57% of the clusters) exhibited overrepresentation of at least one GO term. These 53 clusters amounted to 9,294 IDRs (i.e., 49% of the human IDR-ome), representing extensive overrepresentation of GO terms across clusters of IDRs (**Supplementary Table 2**, **Supplementary** Figure 5, **Supplementary** Figure 6). Although patterns of conservation of bulk molecular features in IDRs were previously observed to be associated with function in yeast (Zarin et al. 2019), we sought to confirm that the widespread overrepresentation of GO terms were not due to bias in manual selection of clusters. To do so, we repeated the analysis using automatically defined clusters and found qualitatively similar overrepresentations to those identified in the manually identified clusters (see **Methods, Supplementary Table 2, Supplementary Table 3, Supplementary** Figure 7). These results indicate a proteome-wide association between the patterns of evolutionary conserved molecular features of IDRs and their biological functions.

To obtain a first global picture of the types of biological functions involving human IDRs, we discerned around functional 20 categories, with some overlap, covering the majority of overrepresented GO terms linked to various IDR clusters (**Supplementary** Figure 6, **Supplementary Table 2**). We defined those categories by grouping related overrepresented GO terms, as detailed in the **Methods**. Some of the most populated clusters are associated with DNA binding (23%), Chromatin/Chromatin binding (33%), RNA metabolism (22%), Cytoskeleton (12%), Signaling (11%), Transmembrane transport (7%), and Reproduction (7%) (**Supplementary** Figure 6), including biological functions that are frequently attributed to IDRs (e.g., DNA binding, RNA metabolism, signaling). We also take note of some less widespread, but significantly overrepresented terms in the IDR-ome map, such as those associated with ‘histone modifications’ (4%), ‘cell morphogenesis’ (2%), ‘innate immune response’ (2%), ‘nuclear pore complex’ (1%) and ‘clathrin binding’ (1%) (**Supplementary** Figure 6). Focusing on overrepresented molecular features in the GO-term enriched clusters (**Supplementary Table 2**), we find that the RNA-associated IDRs are enriched in conserved Arg-Gly/Arg-Gly-Gly (RG/RGG) motifs, Lys (K) content and K homorepeats, Arg (R) and Arg+Tyr (R+Y) content, as well as homorepeats of acidic residues (Asp (D), Glu (E)), all of which is in line with features of IDRs typically associated with phase separation, as many RNA-associated IDRs bind to RNA in the context of biomolecular condensates (Alberti & Dormann 2019; Chong et al. 2018; Youn et al. 2018, 2019). For the IDRs associated with transmembrane transport protein GO terms, we note that many of these IDRs belong to G protein-coupled receptors (GPCRs) (**Supplementary Table 2**). Many of our select clusters show enrichments in biological functions less appreciated to be associated with IDRs (e.g., development, cell morphogenesis, extracellular space, lipase activity) (**Supplementary** Figure 6), suggesting that the diversity of functions in which IDRs are involved is even greater than currently appreciated.

Having established that human IDRs can be clustered based on their evolutionary signatures (**Figure 3**, **Supplementary** Figure 5), and that many of these clusters exhibit enrichments in specific biological functions (**Supplementary Table 2**), we selected a few representative clusters to illustrate this finding. First, as means of validation, we searched for clusters of IDRs with molecular features that have been experimentally confirmed to be critical for specific biological functions. For example, a cluster of 69 IDRs exhibits strong overrepresentation of the GO terms ‘structural constituent of the nuclear pore’ (77-fold, *p* < 1e-9) and ‘signal sequence binding’ (33-fold, *p* < 1e-4) (**Figure 3B**, **Supplementary Table 3 – Tab E**). The nuclear pore complex contains many IDRs and provides binding sites for signal sequence-bearing proteins (e.g., nuclear localization or export sequences) (Mosalaganti et al. 2022; Schmidt & Görlich 2016). The molecular features of the IDRs in this cluster are enriched in specific amino-acid content (Ala, Phe, Gly, Thr), repeats (Gly-Gly (GG), Phe-Gly (FG), Ser-Gly (SG), Pro-Thr-Ser (PTS)), and FG-rich motifs (**Figure 3B**). The specific enrichment in Gly and Phe content and FG repeats is consistent with expectations from the literature, as the IDRs associated with the nuclear pore complex are known as FG-nucleoporins and are rich in Gly, Phe, and FG repeats. For instance, in the essential component of NPCs, FG-NUP98 (Yu et al. 2023a), the amino acids Ala, Phe, Gly, and Thr comprise nearly 40% of its IDRs, where IDRs map to residues 1 to 728, and 888 to 1133. These results confirm that the well-known unusual amino acid distribution and FG-repeats in nucleoporins are conserved during evolution, and that this pattern of conserved bulk molecular features is found in only a small number of IDRs in the human proteome.

A second validation comes from a cluster of 299 IDRs that contains GO-term overrepresentations for ‘mRNA splicing via the spliceosome’ (23-fold, *p* < 0.0001), ‘regulation of mRNA processing’ (19-fold, *p* <0.0001), and ‘positive regulation of mRNA metabolism’ (15-fold, *p* < 0.001) (**Figure 3B**, **Supplementary** Figure 8, **Supplementary Table 3 – Tab F**). These IDRs are not significantly depleted in any feature but have strong enrichments in Gly content, Gly homo-repeats, Arg-Gly (RG) and Arg-Gly-Gly (RGG) repeats, FG repeats, Ser-Gly repeats (SG), and Phe-Gly-Arg (FGR) repeats (**Supplementary** Figure 8, **Supplementary Table 2 – Tab D**). RG and RGG motifs are known to be enriched in IDRs that bind to RNA (Chong et al. 2018) and are implicated in RNA recognition during spliceosome assembly (De Vries et al. 2022). Indeed, biophysical experiments in some RG/RGG-containing proteins have shown that mutation or disruption of RG/RGG motifs can disrupt phase separation and the affinity for RNA (Chong et al. 2018; Ozdilek et al. 2017). These and numerous additional examples (**Supplementary Table 2**) suggest that our clustering of the human IDR-ome based on conservation of molecular features retrieves findings consistent with those reported in the literature.

Next, we examined a cluster of 136 IDRs that is significantly overrepresented in the GO terms ‘histone binding’ (7.7-fold, *p* = 1e-3) and ‘chromatin binding’ (5.2-fold, *p* = 1e-2) (**Figure 3B**, **Supplementary** Figure 8, **Supplementary Table 3 – Tab A**). Histones contain positively charged intrinsically disordered tails that are exposed to solvent (Kim et al. 2023), and these IDRs facilitate interactions, which are often PTM-dependent, with many different histone-binding proteins. Histone-binding domains, such as chromodomains and plant homeodomain (PHD) fingers, often rely on hydrophobic pockets and aromatic residues to bind to methylated lysine residues in histone tails (Li et al. 2006; Nielsen et al. 2002). Interestingly, however, we do not find aromatic residues within the enriched molecular features in our cluster of histone-and chromatin-binding IDRs; instead, we observe strong enrichments in acidic content and repeats of Asp (D), Glu (E), and Lys (K) (**Figure 3B**, **Supplementary Data**). Remarkably, a recent study identified that an acidic TFIIS N-terminal domain (TND)-interacting motif (TIM) in IDRs can mediate binding to histone chaperones and histone readers, including LEDGF and HRP2 (Cermakova et al. 2021, 2023). The TIM motif is characterized by a short α-helix followed by an acidic region, which often contains D or E repeats (Cermakova et al. 2021, 2023). Furthermore, we found numerous clusters with strong overrepresentation for histone demethylase or histone deacetylase activity, which feature conservation of D repeats, Asn (N) residues and N homo-repeats, suggesting how a specific molecular grammar of negatively charged and Asn resides could support DNA binding more generally (**Supplementary Table 2**). We detail functional and feature overrepresentations across clusters in **Supplementary Table 2**.

We then looked for novel functional insights in clusters showing seemingly unrelated enrichments. In the cluster of IDRs associated with the nuclear pore complex (NPC), described above (**Figure 3B**), a significant GO-term overrepresentation was also observed for ‘clathrin binding’ (26-fold, *p* < 1e-5), and the IDRs in the cluster were enriched for SLiMs associated with clathrin-mediated endocytosis (NPF motif) and endosomal sorting complexes required for transport (ESCRT) (**Supplementary** Figure 9, **Supplementary Table 3 – Tab E**). The early stages of clathrin-mediated endocytosis involve many IDRs, including those in NUMB and Epsin-1 that contain the NPF SLiM (Lomoriello et al. 2022). Consistent with this, IDRs from Epsin-1/2/3 (along with PICALM and SCYL2) which are in this cluster, show strong signals for the NPF motif in ELM (EH_1), while most of the FG-NUPs in the cluster do not (**Supplementary** Figure 9). As expected, the IDRs from clathrin-associated proteins do not show conservation of the FG-repeats found in the nuclear pore IDRs (**Supplementary** Figure 9). Given that NPCs are located in the nuclear envelope and clathrin-mediated endocytosis occurs at the cell membrane, the observation of GO-term enrichments in both processes seems unrelated. Functionally, however, the NPC and clathrin-coated vesicles both oligomerize on membrane surfaces and induce membrane curvature (Beck et al. 2018). FG-NUP IDRs and clathrin-associated IDRs cluster together because they all show conserved signals of unusually high hydropathy and a paucity of charged residues (**Supplementary** Figure 9). We speculate that these molecular properties are important for their oligomerization on membrane surfaces. Moreover, key scaffold components of the NPC and clathrin-coated vesicles share similar structural folds and are speculated to have evolved from a common ancestor (Beck et al. 2018; Devos et al. 2004). Recent work has even directly linked NPCs and clathrin-mediated endocytosis: disruption of NPC assembly triggers activation of an ESCRT-dependent chaperone system (Thaller et al. 2019), and clathrin regulates the recruitment of ESCRT proteins (Wenzel et al. 2018). Our observation of similar evolutionary signatures for IDRs that are associated with NPCs and clathrin-mediated endocytosis suggests that evolutionary selection for high hydrophobicity and lack of charged residues may result from similar functionality of these processes.

### “Unexplored” clusters with fully disordered proteins and proteins of unknown functions

The highlighted examples showcase how our IDR-ome map allows the association between biological functions, evolutionary signatures, and specific IDR clusters (**Figure 3**, **Supplementary Table 2**, **Supplementary Table 3**, **Supplementary** Figure 8), thereby establishing a relationship between IDR sequence and function. However, in both automatic and exploratory analysis of clusters, it was evident that several clusters defined by strong patterns of evolutionary signatures did not exhibit any known functional enrichments. For instance, about 43% of the clusters selected in our exploratory analysis do not have any significantly overrepresented GO terms (**Supplementary Table 2**). Some of these clusters may reflect the difficulties and biases in protein annotation (Kustatscher et al. 2022), and we speculate that some of the clusters have no overrepresented GO terms due to missing knowledge of the biological functions of many IDRs and IDPs. For example, the IDR-containing proteins of unknown function ANKRD20A1, FAM131A, and C8orf48 map to “unexplored” clusters that have no GO-term overrepresentation (**Supplementary** Figure 5, **Supplementary Data**). Given the similar evolutionary signatures of the IDRs in the respective “unexplored” clusters, this resource represents an opportunity to discover new biological functions of these IDR-containing proteins. The conserved molecular features of IDRs in clusters without overrepresented GO terms point to properties of these IDRs that are evolutionary conserved, and therefore presumably important for the biological functions of these IDRs. At least four of our selected clusters, based on strong patterns of evolutionary signatures and totaling over 1000 IDRs, show no appreciable overrepresentations of any GO terms.

Next, we investigated where completely disordered proteins, or IDPs, cluster in our map of the IDR-ome, as IDPs are typically challenging to predict biological functions from sequence alone. As compared to IDRs, we found that a higher proportion of IDPs, roughly 31% (226 of 718), are not clustered at all (**Supplementary Table 4)**, indicating that these proteins did not pass one of our filtering criteria (see **Methods**). For the remaining 69% of IDPs (492 of 718) that are clustered, approximately 60% of them (298 of 492) map to clusters with significantly overrepresented GO terms (**Supplementary Table 4**), which is slightly higher than for IDRs in general (49%). Our results provide a valuable resource to further interrogate IDP biology. The IDP-containing clusters that did not yield any overrepresented GO terms, amounting to nearly 40% of the clustered IDPs, thus correspond to “unexplored” clusters.

Some of the IDP-containing “unexplored” clusters contain an abundance of fully disordered proteins (**Supplementary** Figure 10), such as a cluster of 17 IDRs of which 15 are IDPs (88%) from the family of late cornified envelope (LCE) proteins (**Supplementary Table 3 – Tab D**). At first glance, the clustering of different LCE IDPs would appear to be caused by simple positional sequence conservation, as the proteins are all from the same gene family. However, the sequences of these LCE IDPs have highly diverged and traditional bioinformatic approaches reveal no homology between most members (**Supplementary** Figure 10). By contrast, the evolutionary signatures of these 15 different LCE IDPs are all related to one another and show clearly that these IDPs are similar (**Supplementary** Figure 10). Although this cluster does not have any overrepresented GO terms, there are evolutionary signatures from many different molecular features, including strong enrichments in Gly-and Arg-rich motifs (RG, FG, Ser-Gly (SG), Ser-Arg (SR)), dipeptide repeats (Gln-Gln (QQ), Ser-Ser (SS), Gly-Gly (GG)), amino-acid content (Gly, Cys), and SH3-domain binding motif (LIG_SH3_2) (**Supplementary Data**). Another “unexplored” cluster of 21 IDRs with 13 IDPs (62%) shows evidence of conservation for amino-acid content with enrichment in Glu and depletion in Ser residues (**Supplementary Table 3 – Tab B**, **Supplementary Data**). These IDPs are members of the G-, P-, and X-antigen (GAGE, PAGE, XAGE) families, and like the IDP-rich cluster discussed above, show some sequence homology but exhibit far enhanced relatedness when viewed through the lens of evolutionary signatures (**Supplementary** Figure 10). Finally, a cluster of 76 IDRs that contains 42 IDPs (55%) yields strong depletions in amino-acid content (Asn, Asp, His, Leu, Phe) and aliphatic residues, with enrichments in dipeptide repeats (SS, GG), Gly-rich motifs (SG, FG), and Cys content (**Supplementary Table 3 – Tab C**, **Supplementary Data**). These IDPs correspond to 35 different keratin-related (KR) proteins and seven different small glycine and cysteine repeat (SGCR) proteins. Outside of a small group of related sequences, very limited homology is detected by sequence alignments (**Supplementary** Figure 10). By contrast, our evolutionary signature approach shows clear similarities between the 42 IDPs (**Supplementary** Figure 10). Thus, the “unexplored” clusters in our map of the IDR-ome that do not have significantly overrepresented GO terms can provide a unique view of under-characterized IDRs and even fully disordered proteins or IDPs that have similar evolutionary signatures. Furthermore, our analysis of different IDP sequences confirms that our clustering is not caused by positional conservation but is instead due to similar evolutionary signatures.

Finally, we wondered whether human proteins of unknown function that have remained uncharacterized (Duek et al. 2018; Zahn-Zabal et al. 2020) are clustered in our map of the IDR-ome. To this end, we downloaded 1,521 proteins of unknown function from the neXtProt Database (Zahn-Zabal et al. 2020), and we filtered this list to retain only IDR-containing proteins. We found that 58% of the uncharacterized proteins contain IDRs (878 of 1,521 proteins, totaling 2,463 IDRs, **Supplementary Table 5**), which is slightly less than the background proteome (63%). However, over 10% of these uncharacterized proteins are IDPs, which is nearly two-fold higher than the whole proteome. Moreover, although the median IDR length of the uncharacterized proteins was unchanged from the IDR-ome (75 vs. 74 residues, respectively), the median disorder content was significantly increased from 30% in the proteome overall to 53% in uncharacterized proteins (Mann-Whitney test, *p* value < 1e-6) (**Supplementary** Figure 11). Thus, human proteins of unknown function have a higher overall content of disordered residues than the average IDR-containing protein in the proteome (**Supplementary** Figure 11). In our map of the human IDR-ome, over 90% of the 878 IDR-containing proteins of unknown function are clustered (93%; 818 of 878 proteins, **Supplementary Table 5**). This indicates that conserved molecular features within the IDRs of uncharacterized proteins exhibit similarities to other IDR-containing proteins in the proteome and that the sequences of these IDRs pass our quality control criteria (**Methods**). Remarkably, nearly half of the uncharacterized IDR-containing proteins (48%, 396 of 818 proteins) are clustered with significant GO-term enrichments (**Supplementary Table 5**). As an example, we consider the protein of unknown function CDR2 that is predicted to be 92% disordered by SPOTD. The long IDR in CDR2 (residues 1-417) is clustered with 576 other IDRs that are enriched in the Glu-Asp (ED) ratio, Gln content, and helical calmodulin binding motifs (LIG_CAM_IQ_9) (**Supplementary Table 2 – Tab D**). These IDRs are notably depleted in repeats involving Pro, Arg, Ser, Thr, and Gly residues (SS, PR, SG, PTS). The GO terms overrepresented in this cluster are associated with aspects of the cytoskeleton (microtubule motor activity, *p* = 7e-3; tubulin binding, *p* = 1e-4; cytoskeleton organization, *p* = 2e-4) as well as the organization of the cilium (*p* = 1e-2) (**Supplementary Table 2 – Tab D**). Interestingly, a recent affinity-purification proteomics study identified CDR2 as one of the core components in the ciliary interactome (Boldt et al. 2016), with 30 different binding partners identified, including the cytoskeletal proteins tubulin beta chain (TUBB), tubulin beta-4B chain (TUBB4B), and kinectin (KTN1). This example illustrates the potential of our map to link sequence features and biological functions to previously uncharacterized proteins.

Our functional map of the human IDR-ome stratifies human IDRs into clusters that frequently exhibit significant GO enrichments, which helps to reveal the relationship between IDR sequence and function. Our methodology is applicable to fully disordered proteins and proteins of unknown function that are otherwise challenging for conventional bioinformatic approaches. The map is a resource that supports exploration for novel insights by, e.g., comparing clusters with different conserved features of IDR sequences but similar functional enrichments, or identifying groups of IDRs that share conserved features but have no annotated functions yet (see **Code and data availability**). Our analyses above suggest that potential functions can be proposed for a cluster of unknown function once a few of its members have been characterized in depth.

### Specific biological functions can be assigned to human IDRs based on evolutionary signatures

Next, we tested whether a systematic prediction of the biological functions and locations of individual IDRs in the human proteome could be achieved based on evolutionary conserved molecular features. To this end, we applied a machine-learning approach termed FAIDR (Zarin et al 2021) (**Figure 4A**). FAIDR enables the assignment of specific IDRs to specific functions or locations and simultaneously identifies a sparse set of molecular features that are most predictive of (i.e., most associated with) each. The training of FAIDR is based on GO molecular function, biological process and cellular localization annotations, which are available on a per-protein level. If the protein contains multiple IDRs, FAIDR can be used to single out which IDR is most likely involved. The top-scoring predictions for human IDRs, such as ‘nucleolus,’ ‘spliceosomal complex,’ and ‘GPCR activity,’ consistently yield reasonably strong classification performance (AUC values of 0.8 or higher, with corresponding PPV values at 0.4 or above) (**Figure 4A**, **Figure 4B**). Using these AUC(PPV) cutoffs (see **Methods**), we found that we can annotate 148 GO terms to human IDRs (**Figure 4A**, **Supplementary Table 6)**. Many other GO terms we could not reliably predict based on evolutionary signatures, so we do not consider them further here (**Supplementary Table 6**). We found that one of the most strongly predicted GO terms was ‘histone binding’ (AUC = 0.83) (**Figure 4B**). As expected, the positively predictive molecular features of the model align with those overrepresented in the clusters enriched for this term, but these molecular features are distributed across multiple clusters, with alanine content overrepresented in one ‘histone binding’ cluster and di-lysine repeats, along with aspartate-and glutamate-homorepeats, dominating signals in other cluster(s). This underscores the complementary insights that can be derived from an unbiased, unsupervised clustering analysis (**Figure 3**) and the supervised FAIDR approach (**Figure 4**).

**Figure 4.**
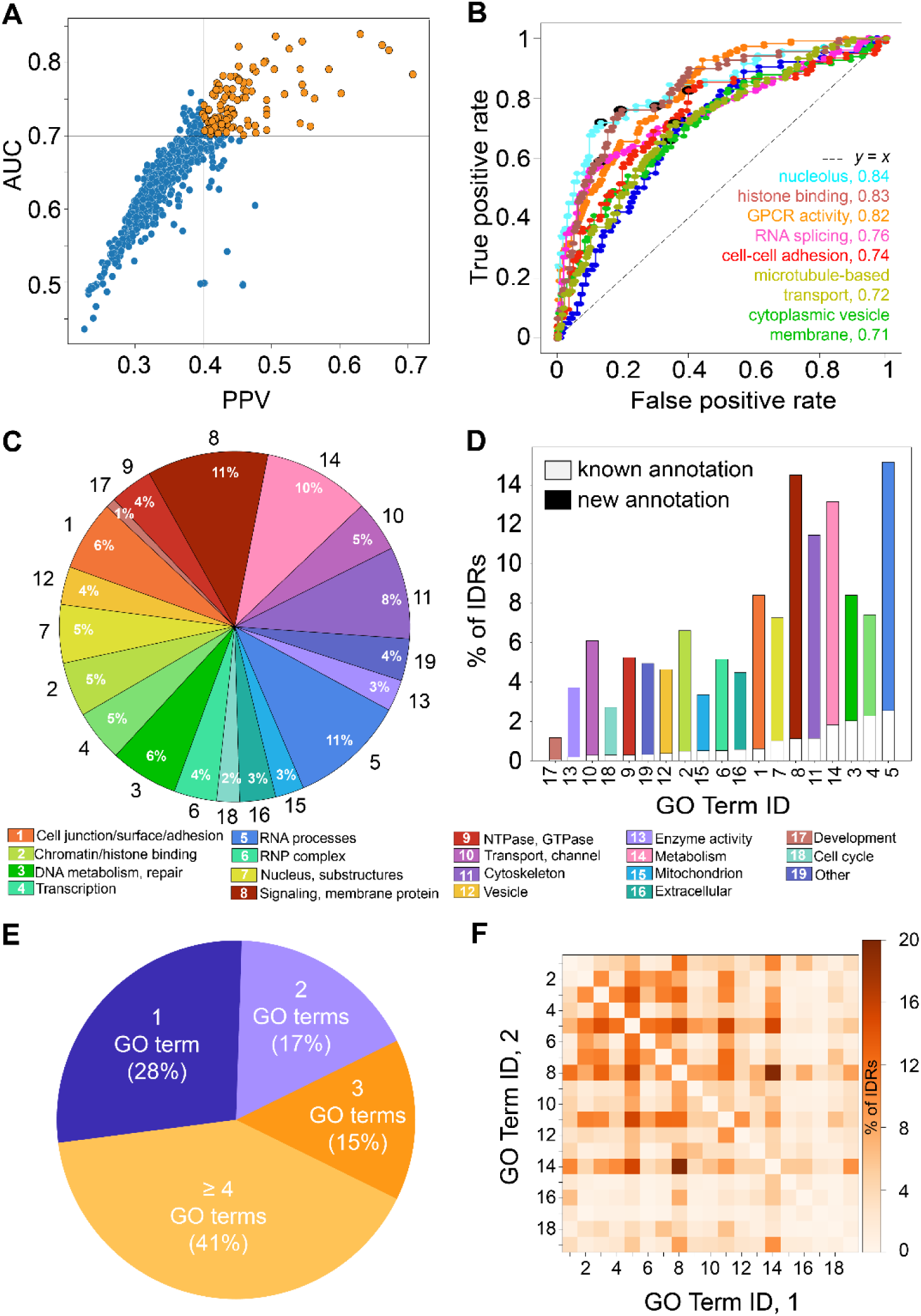
Predicting functions and locations of human IDRs. **(A)** Each point represents held-out data performance of a classifier for one of 600 GO terms covering a broad range of molecular functions, biological processes, and cellular localizations (**Supplementary Table 4**). The X-axis represents positive predictive value (PPV) and the y-axis represents area under the receiver operating curve (AUC) on the held-out data. The models corresponding to data points in orange (PPV > 0.4 and AUC > 0.7) were deemed sufficiently reliable for functional annotations of IDRs proteome-wide (as summarized in **C** and **D**). **(B)** Receiver operating characteristic (ROC) curve for classification of human IDRs to a representative set of GO terms. The performance is shown on held-out data. The terms were selected to display a variation in the FAIDR performance on the binary classification tasks. In addition to the protein level classification, the FAIDR classifier provides probabilities of GO term association for individual IDRs in the protein. These IDR probabilities were used in **C** and **D** to annotate human IDRs to a broad set of enumerated and color-coded functional and location categories. **(C)** A break-up of annotations for all human IDRs to broad functional and location categories as summarized in **Supplementary Table 4** (Tab D). Note that many IDRs get assigned to more than one category (see **E**). In **(D)** the fraction of “known” annotations is given in white boxes. An annotation of an IDR to a function or location was considered as “known” if the protein from which the IDR originates has been previously associated with the category. If the functional or location category was predicted for a protein not previously annotated with that category, and an IDR in the protein was given high probability for the association (see **Methods**), the annotation was considered “new”. See also **Supplementary Figure S12**. **(E)** Each section of the chart represents that fraction of IDRs with a given number of predicted annotations. Only 28% of human IDRs get assigned to a unique functional or location category. **(F)** Correlation between the categories expressed as a fraction of IDRs shared between any given two categories. The diagonal is set to 0. Some of the most frequently shared functional annotations are those associated with various aspects of RNA (5) and DNA metabolism (3 & 4).

Other well-predicted GO terms include “extracellular matrix” (AUC=0.82), “mRNA processing” (0.78), “GTPase regulator activity” (0.77), “centrosome” (0.75), “actin binding” (0.75), “endopeptidase activity” (0.73), and “mitotic cell cycle” (0.70) (**Figure 4A**, **Figure 4B**), once again highlighting the diversity of IDR function in the human proteome (**Figure 4C**). We find that we can ascribe new functions to many more IDRs than were previously annotated (**Figure 4C**, **Figure 4D**), opening venues for hypothesis-driven and validation studies in the future. To provide an overview of FAIDR-based functional annotation proteome-wide, we systematized GO terms into 19 broader, mostly non-overlapping categories, which we refer to as “functional classes” (see **Methods**, **Figure 4C**, **Figure 4D**, **Supplementary Table 6**). For instance, as observed in our map of the IDR-ome clustered by evolutionary signatures, we could ascribe GO terms for biological functions that are often associated with IDRs, including “Transcription” (5%), “Chromatin/histone binding” (5%), and “DNA metabolism, repair” (6%), “RNA processes” (11%), “RNP complex” (4%) and “Signaling” (11%) (**Figure 4C**). However, once again, many IDRs are assigned to GO terms that are not frequently associated with IDRs, including “Cytoskeleton” (8%), “Transport, channel” (5%), and “Cell junction/surface/adhesion” (6%) (**Figure 4C**). Overall, there is a correspondence between the overrepresentations of GO-terms observed in our map of the IDR-ome clustered by evolutionary signatures (**Figure 3**), and those GO-terms we assigned using FAIDR (**Figure 4**).

Through our classification of FAIDR predictions into broader categories, it appears that the majority of human IDRs tend to be associated with more than one functional class (**Figure 4E**). This observation may suggest a certain degree of overlap in our definition of functional classes, but it is also in line with the concept of IDR ‘functional promiscuity’ (Cumberworth et al. 2013). In support of the former, we find the expected overlap in annotations of a broad category “metabolism” (**Figure 4F**, #14) with “DNA metabolism” (**Figure 4F**, #3) and “RNA metabolism, splicing & binding” (**Figure 4F**, #5). However, strong overlaps are also found annotations for “RNA processes (metabolism, splicing & binding)” (#5) and “DNA metabolism” (#3), “DNA binding & transcription” (#4), “signaling (GPCRs, transmembrane)” (#8), and “cytoskeleton” (#11) (**Figure 4F**). Furthermore, predictions for involvement in “signaling (GPCRs, transmembrane)” (#8) are frequently shared with that of “cytoskeleton” (#11). In addition to the discussed overlap with “RNA processes” (#5), we further note signals of shared predictions of “DNA metabolism and repair” (#3) with “signaling” (#8) and of “cytoskeleton” (#11) with “nucleus” and its substructures (“lumen”, “nuclear pore complex”, “nucleolus” and “nucleoplasm”) (#7). Such overlaps in annotations for individual IDRs may point to ‘moonlighting’ (Tompa et al. 2005) or coupling of these biological functions within an IDR sequence, particularly in the case of linking function and localization. Our annotations of functional categories at the IDR level offer a valuable resource for validation studies seeking to unravel sequence-function relationships in greater detail.

To better understand the combinations of molecular features that drive IDR functions, we clustered the FAIDR t-statistic to examine the predictive molecular features for various functions and those that are shared between the most commonly co-occurring functions (**Figure 5**). For instance, among several molecular features that are positively correlated with and predictive of GO terms associated with transcription, we note a high positive Z-score for Pin1 WW domain-binding motifs (DOC_WW_Pin1_4) and for specific SUMOylation motifs, such as KEPE and SUMO-1 (**Figure 5**, **cluster 7**). The role of Pin1 in isomerizing phosphorylated Ser/Thr-Pro bonds in various substrate proteins, including transcription factors and their regulators, has been well established (Hu & Chen 2020; Zhou & Lu 2016). For example, Pin1 binding and isomerization of IDRs can directly regulate the transcriptional activity of key transcription factors, such as STAT3 (Lufei et al. 2007), c-Jun (Wulf et al. 2001), and NF-κB (Ryo et al. 2003). Moreover, among the many proteins that are SUMOylated, transcription factors constitute a large group (Hendriks et al. 2014; Vertegaal 2022), and our data show that evolutionary conserved SUMOylation sites in IDRs are positively predictive of transcription-related GO terms. At the molecular level, SUMOylation of IDRs within transcription factors can regulate activity in various ways, for example by stabilizing a transcriptionally active conformation, such as for SUMOylated HSF1 (Kmiecik et al. 2021), or by controlling subcellular localization, as is the case for NFATC1 where C-terminal SUMOylation induces association with PML bodies (Nayak et al. 2009).

**Figure 5.**
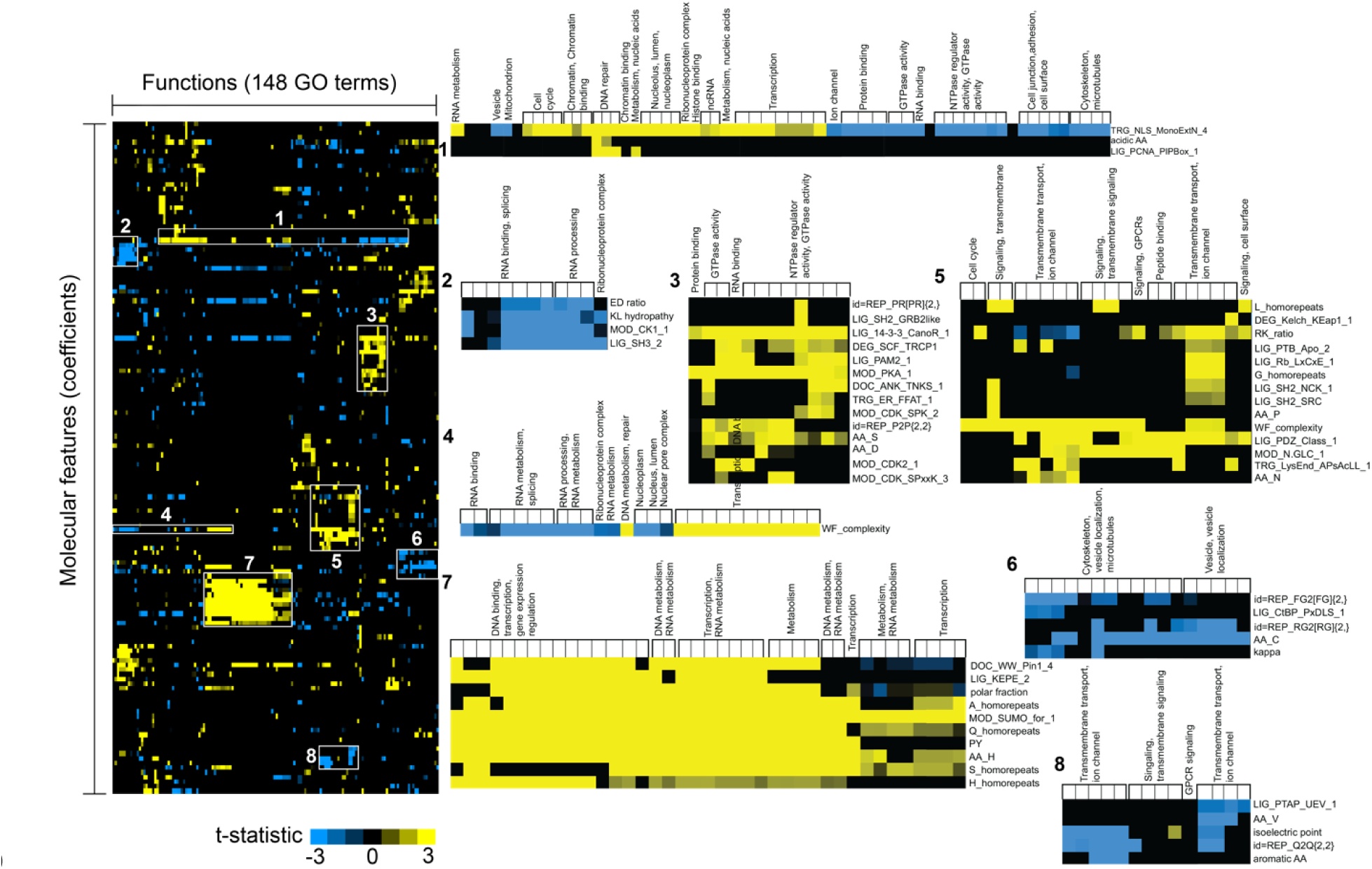
Identifying the molecular features that are predictive of IDR functions and locations. Left heat map represents t-statistics summarizing the predictive importance of different molecular features (y-axis) across GO terms (x-axis). Rows and columns have been organized by clustering. Selected regions, indicated with white rectangles and numbers, are expanded on the right side. Comparisons of the t-statistics within and between different clusters unveils how different combinations of molecular features can drive different biological functions and localization.

Further, our model identifies phosphorylation-related features, including 14-3-3 binding pT/pS motifs and phosphorylation sites for PKA and CDK2, as positively predictive for GO terms that are related to RNA binding and splicing (**Figure 5**, **cluster 3**). The 14-3-3 binding motifs, common in IDRs, are also linked to phase separation of proteins in various signaling pathways (Segal et al. 2023), including those related to RNA processes (Huang et al. 2022). PKA and CDK2 phosphorylation sites have been previously linked to RNA binding and splicing (Ginsberg et al. 2003; Gu et al. 2011; Loyer et al. 2005). Additionally, FAIDR distinguishes the conservation of the PABP-interacting motif PAM2 as positively predictive of RNA binding and splicing (**Figure 5**, **cluster 3**), which confirms the known function of the PAM2 motif in mediating interactions with poly(A)-binding proteins (PABP) and the subsequent recruitment of translation factors and proteins involved in mRNA stability (Kozlov et al. 2010; Xie et al. 2014). Interestingly, we observed that WF complexity is positively predictive for GO terms related to transcription and DNA binding but negatively predictive for terms related to RNA splicing and metabolism (**Figure 5**, **cluster 4**), suggesting that greater sequence complexity in IDRs linked with transcription may hold a specific functional significance. For instance, it is interesting to note that high sequence complexity IDRs linked to transcription or DNA binding often conditionally fold (Wright & Dyson 2015); by contrast, low-complexity IDRs involved in RNA splicing and metabolism are strongly associated with biomolecular condensates (Alberti & Dormann 2019).

We found that phosphorylation-related features, including the PABP-interacting motif PAM2, to be also positively predictive of GTPase activity (**Figure 5**, **cluster 3**). Experimental evidence confirms the association of 14-3-3 proteins with GTPase activity, either by inhibiting it or facilitating the recruitment of proteins in signal transduction pathways (Brandwein & Wang 2017). There are also reports of the involvement of CDK2 and PKA in the signaling pathways of various GTPase families (Ellerbroek et al. 2003; Riou et al. 2013).

Our model also reveals IDR sites that are ligands for PDZ domains and N-glycosylation to be common positively predictive features for GO terms associated with signaling, G protein-coupled receptors (GPCRs), transmembrane transport, and ion channels (**Figure 5**, **cluster 5**). This finding aligns with the recognized roles of PDZ domain-containing proteins as molecular adapters that assemble membrane-associated proteins and signaling molecules (Dunn & Ferguson 2015; Møller et al. 2013; Romero et al. 2011). Additionally, N-glycosylation, prevalent in GPCRs, transmembrane receptors, and voltage-gated ion channels, directly affects signaling via protein-protein interactions, receptor activation, and signal transduction (Goth et al. 2020; Montpetit et al. 2009).

We next compared the FAIDR t-statistic on predicted GO term association for yeast IDRs (Zarin et al 2021) with those of human IDRs. Similar to the Cdc28 targets in yeast, the term ‘cell cycle’ in human IDRs is linked to the presence of CDK consensus and KEN box in the sequence. Both in yeast and human IDRs, the DNA damage response is positively associated not only to the CDK consensus, but also to PIPbox motifs and PIKK (Mec1) motifs. Finally, as we previously noted for yeast IDRs, mitochondrion has a positive association with isoelectric point and aliphatic residues (Zarin et al 2021). These results indicate that the molecular features associated with some biological functions have been preserved over very long evolutionary times.

### The global IDR-ome map reveals positions of disease-associated genes

In addition to the above-described functional insights, the map of the human IDR-ome also reveals positions of genes that are associated with different pathologies. Previously, we reported the enrichment of intrinsic disorder in genes related to complex disease, such as autism-spectrum disorder (ASD) and cancer (Tsang et al. 2020). Here, we asked if our map of the human IDR-ome contains areas where genes associated with these diseases are overrepresented, and if so, what types of conserved sequence features these IDRs contain? We found several clusters of different sizes that contain significant overrepresentation of ASD-risk and cancer genes (**Supplementary Table 7**). At least eight of the clusters contain 10-fold or higher overrepresentation in ASD-risk or cancer-associated genes (*p* values < 0.05, **Supplementary Table 7**), and most of these clusters are not associated with any overrepresented GO terms.

Increased conservation of Q and H residues is evident for several disease gene-containing clusters, with some known to be involved in transcriptional regulation. Interestingly, conservation of the same features was found to be predictive of autism-risk genes by FAIDR (see below). We also note a link between clusters showing enrichment in cancer census genes and stress-granules (**Supplementary Table 7**), indicating a possible association between dysregulated stress granule formation and cancer. Moreover, we also note that some of the IDRs from genes related to ASD cluster together or close to the clusters enriched for cancer genes, which is likely related to shared processes of these IDR-containing proteins, such as transcriptional regulation (Tsang et al. 2020). It is important to note that several clusters enriched for disease-risk genes (both cancer and autism) feature no overrepresentation in any known GO terms (**Supplementary Table 7**). Our results suggest that evolutionary signatures could help understanding of disease genes that encode for proteins with substantial intrinsic disorder.

### Evolutionary signatures enable prediction of localization to specific biomolecular condensates

In cells, many different proteins partition into distinct biomolecular condensates (Banani et al. 2017; Forman-Kay et al. 2022; Lyon et al. 2021). Although the phase-separation propensity of many proteins can be predicted from amino-acid sequences (Cai et al. 2022; Chu et al. 2022; Hadarovich et al. 2023; Vendruscolo & Fuxreiter 2023; Vernon & Forman-Kay 2019), the molecular properties that drive the specificity and composition of different condensates are less well understood.

To evaluate FAIDR’s effectiveness in predicting proteins associated with biomolecular condensates, we utilized two distinct datasets for each condensate category. The training datasets comprised 229, 165, and 519 proteins that were experimentally validated to localize to stress granules, nuclear speckles, and the nucleolus, respectively (Youn et al. 2019, Lu et al. 2019). The test datasets consisted of 32 (cytoplasmic stress granule, GO:0010494), 305 (nuclear speck, GO:0016607), and 234 (nucleolus, GO:0005730) unique proteins annotated for localization to the respective condensates in Gene Ontology Browser (https://www.ebi.ac.uk/QuickGO/). Proteins present in both training and test sets were excluded from the latter and retained in the former. Subsequently, FAIDR was trained on a total dataset of 2,917 proteins for stress granules, 2,224 for nuclear speckles, and 6,454 for the nucleolus. Finally, we assessed the performance of the model proteome-wide, which encompassed a total of 16,115, 16,778 and 12,578 for stress granule, nuclear speckles and the nucleolus, respectively.

We asked whether we could identify the molecular features of human IDRs that lead to their predominant association with a specific condensate. We trained FAIDR on a benchmark of 913 IDR-containing proteins that are known to associate with stress granules (*n* = 229), nuclear speckles (*n* = 165), or nucleoli (*n* = 519) based on experimentally derived datasets (**Methods**). Examining performance across distinct datasets, utilizing Gene Ontology annotations, for 571 IDRs from 553 proteins (**Methods**), revealed that evolutionary signatures of IDRs predict compartmentalization with moderate power (**Figure 6A**), yielding AUC values of 0.69, 0.66, and 0.73 for stress granules, nuclear speckles, and nucleoli, respectively. This performance is comparable to the state of the art, in which AUC values of up to 0.76 were obtained for nuclear punctae proteins (Hadarovich et al. 2023). The top 10% of IDR-containing proteins that we predict to associate with stress granules, nucleoli, and nuclear speckles are provided in **Supplementary Table 7** (Tab E).

**Figure 6.**
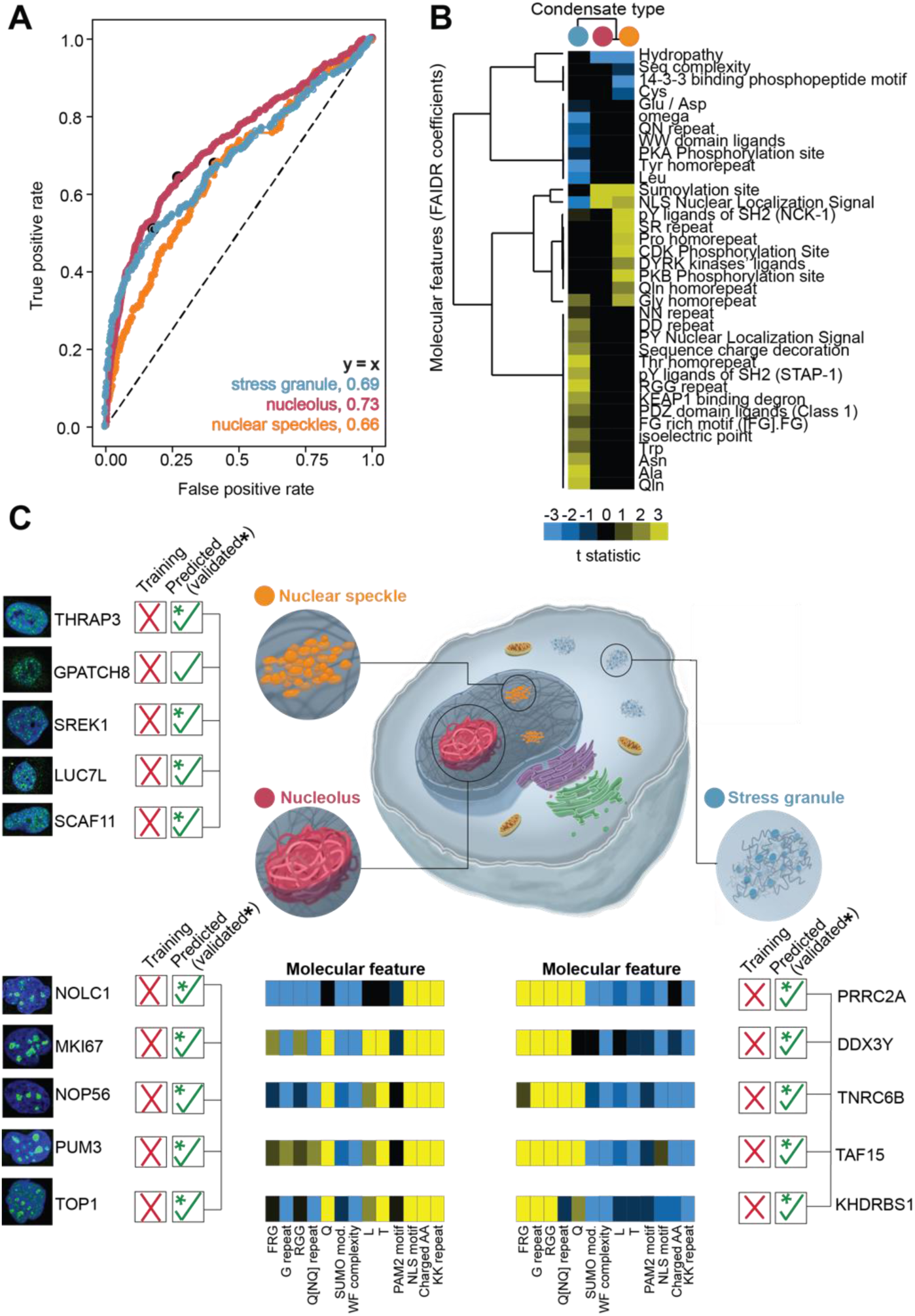
Predictions of association with different bimolecular condensates. **(A)** Receiver operating characteristic (ROC) curve for classification of human IDRs to different cellular bimolecular condensates: stress granules (blue), the nucleolus (red), or nuclear speckles (orange). The performance was tested on independent and non-overlapping datasets with condensate annotations (see **Methods**). Area under the curve (AUC) values are shown in the lower right. **(B)** Hierarchical clustering of the t-statistics as in Figure 5. A negative or positive association of a conserved molecular feature with a particular biomolecular condensate is given in blue or yellow scaling, respectively. The condensate type is shown at the top with the colors the same as panel A. **(C)** Select examples from the top predictions of association to stress granules, the nucleolus, or nuclear speckles. The examples include proteins that were not used in training and for which an association with the indicated condensate was not previously reported. For the nucleolar and stress granule-associating proteins, the evolutionary signatures for selected molecular features are shown with the colors scheme of Figure 3.

We next explored the underlying molecular properties of IDRs that encode the specificity for these different condensates (**Figure 6B**). As an example, consider the IDRs that localize to two different condensates within the nucleus: nuclear speckles and nucleoli. What molecular features of IDRs are responsible for the distinct partitioning into nucleoli *versus* nuclear speckles? As expected, our model identifies that nuclear localization signals (NLS) are important for IDRs that localize to both the nucleolus and nuclear speckle (**Figure 6B**), as well as an increased presence of SUMOylation sites and a decreased overall hydropathy. For nuclear speckles, we find an enrichment in molecular features that include pY ligands of the SH2 domain NCK-1, Ser/Arg repeats, CDK and PKB phosphorylation sites, DYRK kinase ligands, and homo-repeats of Gly, Gln, and Pro (**Figure 6B**). Indeed, nuclear speckle proteins are highly phosphorylated and enriched in Ser/Arg repeats, which have been shown to direct localization to nuclear speckles (Krämer 1996; Li & Bingham 1991). Remarkably, our model correctly identifies Ser/Arg repeats and multiple phosphorylation motifs as distinguishing factors in IDRs that specifically drive localization to the nuclear speckle (**Figure 6B**).

For IDRs that localize to stress granules, the overall pattern of conserved molecular features differs from those of the nuclear condensates (**Figure 6B**). Our model identifies strong enrichments in RGG motifs, FG-rich motifs, PDZ domain ligands, and KEAP1-binding degrons (**Figure 6B**). Multiple sets of experimental evidence confirm that these motifs are abundant in stress granule-containing IDRs (Millar et al. 2023; Youn et al. 2018), such as PRRC2A with several RG and FG, motifs. KEAP1 is an adaptor protein that associates with the E3 ubiquitin ligase CUL3, and the positive predictive value of KEAP1-binding degrons is particularly interesting in the context of recent works reporting on roles of ubiquitylation on stress granule dynamics (Gwon et al. 2021; Maxwell et al. 2021). Other enrichments in bulk properties include isoelectric point; sequence charge decoration; content of Trp, Asn, Ala, and Gln; di-peptide repeats NN and DD; and homo-repeats of Gln, Gly, and Thr (**Figure 6B**). A strong negative signal for the property omega (Martin et al. 2016) suggests selection for well-mixed patterning of charged and Pro residues relative to all other residues (as opposed to blocky patterning) in stress granule-associating IDRs (**Figure 6B**). Interestingly, while a significant depletion in classical NLS is detected for stress granule IDRs, a strong enrichment is found for Pro-Tyr NLS (PY-NLS) (**Figure 6B**), which at first glance appears counter-intuitive with the cytoplasmic localization of stress granules. However, recent work has shown that the stress granule-associated proteins FUS, EWS, and TAF-15, which all harbor PY-NLS motifs that are adjacent to RGG motifs, shuttle between the nucleus and cytoplasm in an Arg methylation-dependent manner (Dormann et al. 2012). Other IDR-rich proteins with PY-NLS signals also undergo nucleocytoplasmic shuttling, including hnRNPA1 and hnRNPA2 that harbor RGG motifs near the PY-NLS (Guo et al. 2018). Thus, evolutionary conserved molecular features within IDRs appear to be sufficiently informative for machine learning protocols to learn aspects of protein specificity for different condensates.

### Leveraging evolutionary signatures to discover condensate-and disease-associated proteins

We next sought to leverage the predictive power of our model to discover new condensate-associated proteins. We filtered our predictions of IDRs associated with the nuclear speckle, nucleolus, and stress granule (**Figure 6C**), and focused on IDRs that were not involved in training. We correctly identify 14 (PPV 70%), 18 (PPV 90%), and 12 (PPV 60%) of the top-20 scoring IDRs associated with the nucleolus, stress granule, and nuclear speckle, respectively. This performance aligns with the recent reports from Hadarovich et al. 2023, where 87.5% of the predictions generated by the PICNIC model were experimentally validated. Experimental evidence in The Human Protein Atlas or elsewhere in the literature provides independent validation of our predicted condensate localization (**Figure 6C**). For example, among the top scoring IDRs in the nuclear speckle are THRAP3, GPATCH8, SREK1, LUC7L and SCAF11, all of which are annotated with nuclear speckle localization by The Human Protein Atlas (**Figure 6C**). For the nucleolus, the IDR-containing proteins NOLC1, MKI67, NOP56, PUM3 and TOP1 are all predicted as nucleolar and validated by literature reports (Ahmad et al. 2012; Chang et al. 2011; Pai et al. 1995; Rallabhandi et al. 2002; Singh et al. 2021). Finally, for the stress granule, FAIDR gives high predictive scores to the proteins PRRC2A, DDX3Y, TNRC6B, TAF15, and KHDRBS1 (**Figure 6C**), all of which are listed as “gold standard” category (tier 1) components of stress granules in the RNA Granule Database (Youn et al. 2019).

We visualized the evolutionary signatures for the top-scoring IDRs predicted to localize to the nucleolus or stress granule (**Figure 6C**, **Supplementary Table 7 – Tab E**). In this representation, we compare the molecular features for individual IDRs that are predicted to localize to the same condensate. For instance, [FR]G motifs in stress granule IDRs are strongly enriched overall (**Figure 6B**) and in four of the five examples in **Figure 6C**. Even though TNRC6B exhibits no evolutionary selection on [FR]G motifs, the remaining molecular features are highly similar to other stress granule-localizing IDRs (**Figure 6C**). Presumably, the absence of evolutionary selection for [FR]G motifs in TNRC6B does not preclude its stress granule localization; instead, it is likely that other molecular properties function in a compensatory manner, *e.g.*, the observed enrichments in G repeats or RGG motifs. Similar trends are seen for the nucleolus-localizing IDRs, where the overall pattern of molecular features in each IDR is similar, even if slight differences exist (e.g., no enrichment in Gln content for NOLC1, **Figure 6C**). As an example, the nucleolar protein GTF2F1 contains a strong enrichment in RGG motifs, which are selected for in stress granule-localizing IDRs (**Figure 6B**) and could hint toward an alternative localization for GTF2F1. Indeed, recent experimental evidence confirms that an interacting partner of GTF2F1, GTF2B, shuttles between the nucleus and stress granules (Qin et al. 2023). Thus, examination of the molecular signatures for individual IDRs can provide additional insight into the molecular properties and localization of these IDR-containing proteins.

Finally, the significant overrepresentation of genes encoding long IDRs in ASD risk, as documented by (Tsang et al. 2020), along with the observed enrichment of ASD-risk genes in various regions of the IDR-ome map, implies that conserved characteristics of IDRs may offer valuable insights into this complex disease. We wondered if using FAIDR solely with evolutionary signatures of IDRs could adequately predict ASD-risk genes. To this end, we trained FAIDR using a curated set of IDRs from ASD-risk genes identified by (Satterstrom et al. 2020), and applied the model to estimate ASD-risk proteome-wide (**Methods**). Through leave-one-out validation, we achieved a retrieval rate of 34% for known ASD-risk genes, with an accuracy of 0.6 and precision of 0.4. Notably, among the top 10% of predicted risk genes across the proteome, we identified several genes newly added to the Simons Foundation Autism Research Initiative (SFARI) database in 2023 (Abrahams et al. 2013) (**Supplementary Table 7**). These and other novel predictions indicate that a straightforward model based on IDR features could offer predictive power for identifying new ASD genes.

## Discussion

We measured evolutionary conservation of nearly 150 bulk molecular features (Zarin et al. 2019) (**Supplementary Table 1**), including motif and repeat content and diverse physicochemical properties, in nearly 20,000 IDRs within the human proteome. We show, through both clustering (**Figure 3**, **Supplementary** Figure 5) and classification (**Figure 4**, **Figure 5**, **Figure 6**) analysis, that combinations of these molecular features are associated with diverse biological functions and localizations. Hence, by recasting the sequences of IDRs into evolutionary conserved molecular features, we found a way to connect human IDR sequences to function without relying on multiple sequence alignments, which are not usually observable for IDRs. Our results lend further support to the idea that selection for or against specific features suggests a link between biological function and IDR sequence, as previously established for budding yeast and *Drosophila* IDRs (Singleton & Eisen 2023; Zarin et al. 2019). Indeed, our feature-based concept has been increasingly recognized and used to gain further insight into function and localization of IDR-containing proteins (Cohan et al. 2022; Duffy et al. 2022; King et al. 2024; Loureiro et al. 2021; Millar et al. 2023).

Using the patterns of conservation in IDRs, we established a “map” of the human IDR-ome, in which we can explore groups of IDRs with similar function or location. The map of the human IDR-ome introduced here represents a resource for discovery of functional elements for vast parts of the human proteome, which have thus far eluded standard bioinformatic approaches. We find that the map of the human IDR-ome recapitulates some known biological functions or processes mediated by IDR-containing proteins, such as overrepresentation of GO terms related to DNA-and RNA-binding, but also sheds light on new or under-appreciated functions of IDRs, including their involvement in development and transmembrane transport. For around 40% of the clusters, there are no known GO term annotations, which likely reflects some of the biases and difficulties associated with functional annotation (Kustatscher et al. 2022), particularly for proteins having a large fraction of disordered residues. Importantly, the “unexplored” clusters of IDRs with similar conserved molecular features but no known function are prime candidates for discovering new biology. For instance, a limited set of IDR-containing proteins would need to be examined in order to assign plausible functions to other IDRs in the cluster. Together, our work provides, to our knowledge, the first comprehensive functional map of the human IDR-ome based on evolutionary signatures. The map reveals known and novel combinations of specific molecular features that drive the rich complexity and promiscuous nature of IDR functions.

Our initial functional map of the human IDRs stands to be improved in several ways. First, it is based on a curated list of molecular features that is limited. The list of relevant molecular features will likely increase in the future, and efforts have already been taken to discover functionally relevant features in a systematic and unbiased way using self-supervised deep learning approaches (Lu et al. 2022). Second, we used a combination of unsupervised (clustering) and supervised (classification using FAIDR) analyses to make predictions about IDR function. In part, we rely on this two-stage approach because the numbers of IDRs with some known function (such as those found in the nuclear pore, **Figure 3**) is too small (*n* = 64) to train a standard supervised classifier in a space of nearly 150 features. Future approaches such as semi-supervised, transfer-learning or data augmentation (Lee et al. 2023; Lindorff-Larsen & Kragelund 2021; Lu et al. 2022; Pang & Liu 2023) approaches will likely address these challenges.

Understanding the impact of disease mutations in IDRs is a key area of research. Outside of IDRs with strong positional alignments, which often conditionally fold (Alderson et al. 2023; Piovesan et al. 2022), it is challenging to interpret disease-associated mutations that map to IDRs, which have no stable tertiary structure and, by corollary, limited positional sequence conservation. Here we looked at overrepresentation of genes involved in two diseases in which IDRs feature prominently, autism spectrum disorder (ASD) and cancer (Tsang et al. 2020). The map of the human IDR-ome reveals specific clusters that show significant enrichments in ASD-risk and cancer census genes. Based on these results, we hypothesize that mutations that disrupt conserved features of IDRs in those clusters are more likely to have a pathological impact, a focus of our future research. Functional prediction within IDRs at the residue level is a rapidly growing research area (Barik et al. 2020; Hu et al. 2021). However, these efforts focused on relatively few broad functions, such as protein binding, DNA binding, RNA binding, and linker or ‘entropic chain’, ‘assembler’, ‘scavenger’, ‘effector’, ‘display site’, ‘chaperone’ (Pang & Liu 2022), Residue-level prediction approaches that can more closely approach the diversity of IDR function we observed in the proteome will likely improve the resolution of the initial map presented here, leading to insight into the functional impact of disease mutations.

Although certain IDRs cluster together and are associated with equivalent biological functions, the clusters alone do not precisely inform on specific aspects of IDR function. Which of the evolutionary conserved molecular features are responsible for function? In full-length proteins with multiple IDRs, do one or more of the IDR participate in the biological function? To answer these questions, we used FAIDR to predict association with 148 different GO terms, stratified into 19 functional categories, across the human IDR-ome (**Figure 4**, **Supplementary Table 6**). Our predictions reflect the rich complexity of IDR-driven functions, suggesting that over 60% of IDRs could be assigned to more than one functional category. Here, however, we caution that even a modest rate of false positives would impact the number of the predicted functions. We therefore prefer to consider these predictions in a qualitative way and use them to discover and contrast features strongly associated with different IDR functional categories (**Figure 5**).

An active area of IDR research focuses on the role of particular IDRs in phase separation and formation of biomolecular condensates (Borcherds et al. 2021; Rostam et al. 2023). Although methods exist to predict the phase-separation propensity of an IDR from its amino-acid sequence (Cai et al. 2022; Chu et al. 2022; Hadarovich et al. 2023; Vendruscolo & Fuxreiter 2023; Vernon & Forman-Kay 2019), it remains challenging to understand how condensates achieve specificity and why certain IDRs localize to certain condensates. We showed that FAIDR can not only reliably predict which IDRs will localize to the nucleolus, nuclear speckle, or stress granule (AUC values of *ca.* 0.7 on independent test sets), but can also reveal which conserved molecular features are responsible for specificity. Our performance, with AUC values from 0.66 to 0.73, is near the state-of-the-art (AUC range 0.52-0.76) reported from four different algorithms when tested on a larger set of proteins that localize to nuclear punctae (Hadarovich et al. 2023). A particularly striking example of our feature-based approach is provided by the protein GTF2F1, which is predicted to localize to the nucleolus and whose molecular features therefore resemble other nucleolar-predicted IDRs. However, GTF2F1 contains a significant enrichment in conservation of the molecular feature ‘RGG motifs’, which is otherwise associated with stress granule-localizing IDRs. The RNA Granule Database lists GTF2F1 as a low-confidence potential stress granule protein (tier 4), which could suggest that GTF2F1 shuttles between nucleoli and stress granules depending on specific cellular conditions. While localization by binding of folded domains to specific targets likely contributes, considering the evolutionary conserved molecular features of IDRs can provide key insights into the complex interactions that underlie the specificity of biomolecular condensates, as well as possible modes of regulation.

Finally, we note that widespread association of bulk molecular properties in IDRs with diverse biological functions suggests that the evolutionary characterization of IDRs could be expanded in scope with additional biophysical properties. For example, recent efforts to characterize the structural ensembles of the human IDR-ome using coarse-grained molecular dynamics simulations found some association between chain compaction and biological function (Lotthammer et al. 2023; Tesei et al. 2023). Here, by considering the evolutionary conservation of sequence-based molecular features, we find strong and wide-ranging functional association for nearly 50% of human IDRs. Based on our results, we anticipate that placing IDRs within a higher-dimensional evolutionary and biophysical space will be a key step toward an improved understanding of the molecular basis for cellular function of intrinsically disordered protein regions (Holehouse & Kragelund 2023).

## Methods

### Prediction of intrinsic disorder and boundaries definition

The reference human proteome assembly was downloaded from UniProt (Proteome: UP000005640) in August 2019. We note that “miniprotein” products of short open reading frames, many of which are likely to contain IDRs, are increasingly recognized as functionally important constituents of the human proteome (Duffy et al. 2022). However, such “miniproteins” are not yet included in the reference proteome and were thus not considered here. SPOT-Disorder predictor v1.0 (Hanson et al. 2017) was used to predict the per-residue probability of intrinsic disorder for every protein sequence in the human proteome. We used SPOT-Disorder v1.0 because it provided the closest agreement with NMR-determined disordered content (Dass et al. 2020; Nielsen & Mulder 2019) and is among the most accurate predictors overall (Necci et al. 2021). A disorder probability above 0.5 was used to define disordered residues. Only protein regions with 30 or more consecutive residues that were predicted to be intrinsically disordered were considered as IDRs in all subsequent analyses.

### Computation of IDR fractions and length distribution

To compute the distribution of IDRs in the human proteome, we used the SPOT-Disorder predictions as described above to identify IDRs of 30 or more consecutive residues. Any protein without an IDR was classified as a folded protein. Fully disordered proteins were defined as containing 95% or more disordered residues. Mixed proteins, which contain both IDRs and folded domains, were filtered to test if IDRs are more likely to appear as terminal regions or as linkers that are interspersed between folded domains. We mapped folded domains to each protein via the PFAM database (Mistry et al. 2021) and then determined the relative location of the IDR(s). The histogram of IDR lengths was fit to a power-law distribution (*y* = *Ax^b^* + c) using the scipy package in Python. The fitted parameters and the associated errors derived from the covariance matrix are: *A* = 7.87 ± 0.28 x 10^5^, *b* = -1.55 ± 0.01, *c* = -9.38 ± 1.91.

### Computation of positional sequence conservation in IDRs

Positional sequence conservation was computed on alignments of human IDRs to IDR sequences from orthologous species (see *Retrieval of orthologous protein sequences*). We computed positional conservation across MSA columns using a modified metric of Shannon’s entropy, the so-called property entropy as previously introduced by Capra and Singh (Capra & Singh 2007). Gaps were ignored in the computation of positional conservation. As the gap content of IDRs is relatively high compared to folded domains, and the metric considers only the alignable columns, it grossly overestimates the positional conservation of IDRs.

### Computation of sequence similarity between IDRs in the IDRome

We ran BLASTP (Altschul et al. 1990) to determine to which extent human IDR sequences have positional similarities with one another. BLASTP was run with E-value cut-off values of 0.0001. Any hit within the threshold (other than to query) was considered homologous. For the IDP sequence analysis (**Supplementary** Figure 10), for each IDP cluster we constructed a BLASTP sequence library with the corresponding IDP sequences. We then subjected each IDP in the cluster to a BLASTP search. For any hit with an alignment coverage of at least 50% of the query sequence length with at least 30% sequence identity, we computed the product of these two values (alignment coverage * sequence identity) as a proxy metric to reflect the degree of positional sequence similarity. We ignored any alignment gaps. As a numerical example, an alignment of two sequences with 100% (50%) coverage of the query sequence length and 80% (30%) sequence identity would yield a value of 0.80 (0.15). For any IDP sequence whose alignment to another IDP did not meet the above two criteria, or was not reported in the BLASTP output file, we assigned a value of zero. We then plotted these data as a heat map, and we imposed symmetry on the heat map by taking the larger value between (*i,j*) and (*j*,*i*) and setting this as the value for both coordinates in the final plot. The range of values is between 0 and 1, and thus reflects the similarity of IDP sequences in the respective cluster as measured by a conventional sequence alignment approach (**Supplementary** Figure 10). For the corresponding plot based on evolutionary signatures of these IDPs, we computed the cosine similarity metric, defined as the dot product of the two vectors divided by the product of the norms, between each IDP sequence in the cluster and all other IDP sequences in the same cluster. The input vectors were the 144-dimensional evolutionary signatures. In this representation, IDP sequences with similar (different) evolutionary signatures have values closer to 1 (0) (**Supplementary** Figure 10). For both the alignment and evolutionary signature representations, the matrices are sorted alphabetically by UniProt ID on the *x* and *y* axes.

### Molecular features definition and computation

The majority of the molecular features used in this study were previously defined and summarized in Supplementary Table S4 of the work by (Zarin et al. 2019). Definitions and details of computation of the additional features introduced in this study are listed in **Supplementary Table 1**, which also documents the features that have been modified or adapted for calculation speed purposes. For example, we updated the list of SLiMs (Kumar et al. 2022), made it compatible with the human proteome, and added additional features that have been reported as important for human IDRs in the recent literature (**Supplementary Table 1**) (Banani et al. 2017; Bremer et al. 2022; Chavali et al. 2017, 2020; Kuechler et al. 2020; Lyon et al. 2021; Martin et al. 2020).

We introduced several methodological changes to work with IDRs in the human proteome. First, the human proteome is roughly five times larger than the yeast proteome and significantly more complex, which is reflected in imperfect annotations of orthologous and paralogous genes to human genes in other species. Second, it is difficult and time-consuming to obtain reliable inferences of phylogenetic trees of metazoan species. Hence, the elaborate protocol for null hypothesis computation based on inference of evolutionary distances from a phylogenetic tree outlined by (Zarin et al. 2019) constitutes a bottleneck for a fast and efficient computation of human IDR evolutionary signatures. Therefore, we resorted to a substantially simplified protocol, as described in the sections below. The modified protocol provided similar results as the original method (**Supplementary** Figure 3, **Supplementary** Figure 4).

### Retrieval of orthologous protein sequences

The ‘Retrieve/ID mapping tool’ from UniProt was used to obtain Ensembl gene identifiers for every UniProt identifier that is associated with each canonical protein sequence of the human proteome. The Ensembl gene identifier was used in a call to the Ensembl API (Method: GET homology/id/:id) to dynamically access all orthologous protein sequences that map to the given Ensembl gene identifier. The retrieved sequences were subsequently filtered to keep only those that were annotated as orthologous to the human Ensemble protein identifier and had the amino-acid sequence that matched to the UniProt query sequence. The check for the exact match to the canonical UniProt protein sequence was necessary, as one Ensembl gene identifier matches to several different Ensembl protein identifiers, and annotations based on identifiers alone (UniProt ID to Ensembl protein ID) may not always be accurate. Beyond this check, we relied on the Ensembl annotation of orthology to the canonical human protein sequence.

### Defining orthologous IDRs and computing evolutionary distance

Following the retrieval of orthologous protein sequences, sequence alignments were computed using MAFFT (Katoh & Standley 2013), with the human protein sequence used as a reference. The IDR regions from orthologous sequences were extracted based on the sequence alignments and using the IDR boundaries as defined for the human sequence, as performed previously for yeast proteins (Zarin et al. 2019). For each orthologous IDR sequence, an estimate of a pairwise evolutionary distance to the human IDR sequence was inferred. To this end, we used the Felsenstein 1981 model (F81) (Felsenstein 1981) model applied to proteins (Lemey et al. 2009). Under this model, we can compute the probability, *p*, that any two sites are different at scaled evolutionary distance, *d*, in substitutions per site:

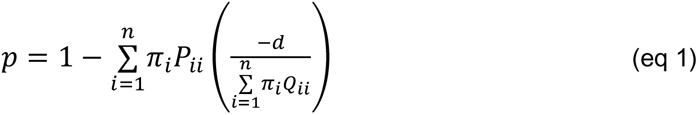

where *πi* are the amino-acid probabilities in the extant human IDR sequences, *i* indexes the alphabet of 20 amino acids, and the 20 x 20 substitution rate matrix *P* is defined as in the F81 model (Felsenstein 1981) applied to proteins:

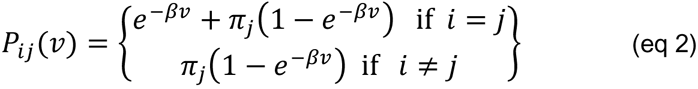

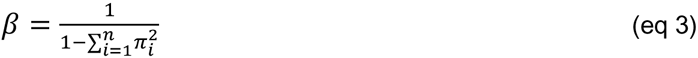

where *v* represents the expected number of amino-acid changes per site. *β* can then computed as given in *eq 3*. To estimate *p*, the probability that two sites are different, we simply divide the total number of different positions between the human IDR and an orthologous IDR by the total number of aligned positions, so that gapped positions (i.e., indels) were ignored in the enumeration of *p*.

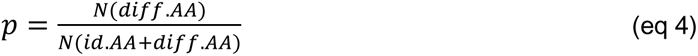

Given the fraction of positions that differ between the reference human IDR and the query orthologous IDR (our estimate of *p*), and *β* as defined above, the estimate of the evolutionary distance *d* is obtained by substituting and rearranging eq. 1 as:

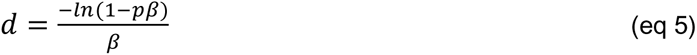

Note that this evolutionary distance (*d*) is a method of moments estimator (Lemey et al. 2009). We highlight that this simplified method does not require inference of a phylogenetic tree, or any other information from the alignment other than the proportion of sites with different amino-acid identities between the reference (human) and the query orthologous sequence.

### Orthologous sequence selection

To select a set of orthologous sequences that give a wide distribution of approximate evolutionary distances (see above) to the corresponding human IDR sequence, we employed an iterative heuristic. The human IDR sequence was used as a reference in all comparisons. In the first steps, any orthologous sequences that were *‘length_factor’*-fold too short or too long relative to the reference were discarded. A *length_factor* of 3 was used in all comparisons. On the remaining sequences, we first calculated a pairwise distance from the reference using the approach described above. The distance *d* was computed as given in *eq 5,* with *β* computed from amino acid frequencies in the human IDR-ome as given in *eq 3.* Distances were capped at a value of 10 in instances where no amino-acid matches were found between the two compared sequences, and when all amino-acid positions were equivalent the distances were set to 0. Any sequences with too large (10) or too small (0) a distance from the human sequence were automatically discarded. To obtain a sufficiently diverse set of orthologous sequences, we next employed a heuristic that selects for orthologous sequences that are sufficiently distant from the reference human sequence and from one another. Two parameters control the heuristic: 1.) *d_ratio*, the ratio of distance from the closest neighbor in approximate evolutionary distance space relative to the reference, and 2.) *d_total*, the total sum of approximate evolutionary distances of orthologous IDR sequences from the reference. To select for sufficient divergence and spread, only the sequences that are at least *d_ratio* further away from the reference than from the closest distance neighbour were kept. Furthermore, the sequences were collected until the maximum of the total sum of distances (*d_total*) from the reference was reached. Setting *d_ratio* to a smaller value imposes a strict restraint on the spread of orthologous sequences from one another in approximate evolutionary distance space. On the other hand, setting *d_total* to a smaller value imposes a stricter restraint on the tolerated divergence of the sequences from the reference. We used *d_ratio* of 5 and *d_total* of 30 in all comparisons. Approximately 2,000 human IDRs were removed from our analysis due to a lack of orthologous sequences or too high or too low sequence similarity among orthologs, amounting to a total of 19,032 human IDRs on which our protocol could be applied.

### Simulated null expectation

We substantially changed simulations of null expectation relative to the approach originally applied to yeast (Zarin et al. 2019), as detailed below. In addition, Zarin *et al*. preserved positionally conserved short linear motifs (SLiMs) in simulations of null expectations, which led to an underestimate of the evolutionary conservation of SLiMs. Here, we removed any such conservation requirements for SLiMs. Our simulation of the null expectation was based on the estimated pairwise evolutionary distances between the human IDR sequence and each orthologous IDR sequence that met the specified quality criteria (see *Orthologous sequence selection*). Substitutions and insertions and deletions were simulated independently and combined at the end.

First, given an estimated distance of an orthologous IDR sequence from the reference human sequence *d*, and *π_i_* and *β* (defined by amino-acid frequencies in the extant human IDR sequences, *eq 3*), a substitution matrix *P* was computed based on the F81 model (Felsenstein 1981) as described in *eq* 2. For each orthologous IDR sequence, a set of 1000 simulated sequences were computed from the reference human IDR by amino-acid substitution at each position. The rate of amino-acid substitutions (*i->j*) were given by the probabilities as defined by *P (eq 2)*. The simulated sequences initially all had lengths that were equal to the reference human IDR.

In the next step, insertions and deletions (henceforth indels) were added to each sequence following a Poisson process. Given *d,* the indel rate (*R*_indel_) was defined as:

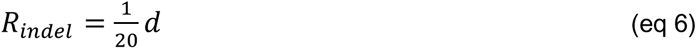

A Poisson process was used to get the positions in the sequence at which an indel occurs given *λ=R_indel_* length_seq.* At every indel position in the sequence, the indel type was assigned at random to either ‘insertion’ or ‘deletion’ with equal probability. The size (*k*) of an indel at position was assigned from an empirically derived distribution:

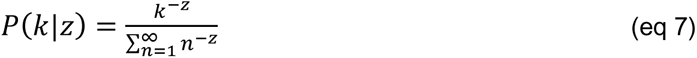

where *z* = 1.5, as previously established (Cartwright 2006; Nguyen Ba et al. 2012). Residues within the inserted segments were drawn according to the amino acid probabilities found in human disordered regions (*π_i_*).

### Computation of evolutionary Z-scores

The molecular features were computed for every human IDR sequence, as well as each of the selected orthologous IDR sequences. In total, 144 different features were computed for each sequence (see above, *Molecular features definition and computation*). The features were also computed for each sequence of the sets of 1000 simulated IDR sequences. The mean Z-score of a feature was obtained by taking the difference between the observed mean of the feature for the selected orthologous IDRs (*x*) and the mean of the simulated means, i.e., the mean of 1000 means from the simulated distributions of the feature, *(μ*), normalized by the standard deviation of the simulated means (*σ*), as introduced previously by Zarin (Zarin et al. 2019).

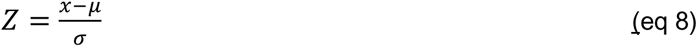

### Z-scores post-processing and clustering

The computed Z-scores measure evolutionary conservation of the molecular features. Prior to any clustering, we removed any molecular features that yielded no Z-scores for any of human IDRs. The inability to compute a Z-score arises when a feature is absent in IDRs of the selected extant species, or due to numeric issues when computing a standard deviation under the null hypothesis for features that are rare and difficult to generate with a simulated null model (e.g., typically SLiMs with long regular expressions and some rare homorepeats, *e.g.*, Trp homorepeat).

Next, to hierarchically cluster evolutionary Z-scores for all human IDRs we used Cluster3.0 (de Hoon et al. 2004). We used the Cluster3.0 interface to first filter the data to remove any entries, that is IDRs, which did not have any significant Z-scores, i.e., Z-scores with absolute value of 3 or more. We then proceeded with default settings for “Hierarchical” clustering using uncentered correlation distance as a similarity metric and average linkage as a clustering method. We calculated weights and clustered IDRs (“Genes” in Cluster3.0 interface). The clustering output was visualized using JavaTreeview (Saldanha 2004), which allows for interactive exploration and export of clusters of interest.

To select the clusters for further analyses, we relied on both manual selection from the displayed hierarchical tree calculated by Cluster3.0 (e.g., **Figure 3**), as well as on an automated selection of clusters given a fixed distance. The distance refers to the uncentered correlation distance between the vectors representing evolutionary Z-score of human IDRs, which is the same distance used in the hierarchical clustering by Cluster3.0. The automated selection was performed on a range of clustering thresholds from *d* = 0.1 to *d* = 0.9 in increment steps of 0.1 (**Supplementary Table 3**). The automatic approach enables a more rapid analysis of data following the initial clustering step and any subsequent re-clustering using a clustering method of choice.

Details of the GO analyses on the clusters and definitions of “broader functional categories” are given below (see *Gene ontology overrepresentation, Definition of functional categories*). To confirm that the functional overrepresentations that we identified were not the result of our potential bias in the selection of clusters, we also tested proteins extracted from automatic clusters for GO overrepresentation (**Supplementary Table 2**). Because it can be performed at a range of distance thresholds, the automatic analysis often reveals functional and feature enrichments that might elude the limited exploratory manual analysis of the IDR-ome map (**Supplementary Table 2**). Regardless of the distance threshold used to define and extract the automatic clusters, a minimum of 30% of the clusters show significantly overrepresented GO terms for molecular function, cellular component, or biological process (**Supplementary Table 2**). This suggests that evolutionary conserved molecular features can be used to effectively cluster the human IDR-ome and reveal functionally related IDRs. A comprehensive analysis of selected clusters using both exploratory analysis and automatic clustering, GO term overrepresentation, and enrichment in positive or negative Z-scores of molecular features relative to the background of the entire IDR-ome is given in **Supplementary Table 2**.

## FAIDR

The FAIDR model was used as published in its original version (Zarin et al. 2021), with adaptations to support training models iteratively on sets of different functional categories and testing each model on independent test sets. FAIDR was applied to our set of 19,032 IDRs from 11,640 unique full-length proteins, of which 8,353 had annotated GO terms. The GO annotations on the protein (i.e., gene) level and the associated evolutionary signatures of the IDRs were used as training data for FAIDR. The GO terms were obtained from the PANTHER database (Mi et al. 2010) (release PANTHER18.0, downloaded in October 2022). We built separate FAIDR models for each of the 601 GO terms listed in **Supplementary Table 6**. To comprehensively test the performance of the FAIDR models, for each model we took bootstrap test samples from the available data and performed model training on the remainder of the data. This allowed us to assess the performance on the entire dataset, whilst ensuring that the performance was tested on examples that had not been seen in training (**Supplementary Table 6, Methods).** We defined a threshold for a reliable FAIDR prediction as a model that on the test set had an AUC (area under the receiver operating characteristic curve) value above or equal to 0.7 and PPV (positive predictive value) above or equal to 0.4 (**Figure 6A**).

### Cross-validation of FAIDR models and functional annotations on the proteome level

Having established that we can successfully identify which IDRs and which molecular features are associated with different gene ontology (GO) terms, we next employed FAIDR to make functional annotations for all IDRs in the human IDR-ome (**Figure 6C**). To this end, we performed cross-validation of 601 FAIDR models trained on 600 GO terms, obtained from PANTHER as mentioned above. For training of FAIDR models, GO term annotations on gene (i.e., protein level) were used. These were assigned as given in the PANTHER database. Upon training, FAIDR models were used to compute the probability of association of each IDR in the proteome with each of the GO terms (Zarin et al. 2021). Only the models trained on a minimum of 100 distinct instances were considered.

For each GO term (*i.e.*, each FAIDR model), the full dataset consisted of 19,032 IDRs. These IDRs were divided into training and test sets six times using sampling with replacement. In each of the six iterations we held out approximately 15% of the IDRs from the positive class (class 1), which had not yet been included in any of the previous test sets. The negative class in each iteration varied as it was set to be three times bigger than the current positive set. This ensured that the positive class of test sets in each iteration were distinct and representative of different portions of the full dataset. After evaluating FAIDR’s performance on the current test set using ROC analysis, we reintegrated it back into the dataset. Consequently, in the next iteration, a different 15% of the positive class of the full dataset was chosen as the test set, which allowed for comprehensive cross-validation of FAIDR across the entire proteome. This approach ensured a robust evaluation of the model’s effectiveness and minimized potential biases. Cumulative ROC statistic of all six iterations was used to establish a threshold for performance of each FAIDR model. For functional annotations of human IDRs across the proteome, only the models for GO terms that achieved an AUC of 0.7 or higher and a PPV of 0.4 or higher were used, which amounted to 148 GO terms.

To assign human IDRs to each of 148 GO functional terms, we relied on likelihoods computed by our regression model for each IDR (Zarin et al. 2021). Based on the receiver-operating-characteristic (ROC) analysis from testing of each model on held-out data, we defined the threshold value of likelihood that minimized false positives while maximizing true positives (**Supplementary Table 6**). To further minimize the chance of false positive annotations, we only accepted an annotation of a function for a protein if the computed probability for the protein was larger than or equal to twice the optimal threshold likelihood. In addition, we only annotated IDRs to GO functional terms if computed probability exceeded 0.6 in case of one or two IDRs in a protein and 0.5 in case of more than two IDRs in a protein. The final sets of functional annotations were pruned to remove redundancies stemming from similar performances of FAIDR models on closely related GO terms. Finally, the annotated GO terms were aggregated to a set of 19 broader functional categories, as summarized in **Supplementary Table 6**. An annotation of an IDR to a category was considered as “known” if the protein from which the IDR originates has been previously associated with the category. If the category was assigned *de novo* to a protein and an IDR in the protein was given high probability for the function (see above), the annotation was considered “new”. The 19 broader functional categories (**Figure 6**) were defined as detailed below. In instances of multiple annotations, pseudo counts were added to the total number of IDRs to compute the displayed percentages.

### Gene ontology overrepresentation

As mentioned in the FAIDR section, the GO term annotations were extracted from the PANTHER database (Mi et al. 2010). To compute GO term overrepresentations for IDRs in select clusters, we used GO annotations available on the protein (gene) level. We considered a set of proteins (genes) in a cluster, i.e., we did not count any duplicates of proteins that might have more than one IDR in the same cluster. To compute overrepresentation, the set of proteins from a cluster was compared against a background of the human standard proteome set, filtered to include only the proteins that contain long IDRs (defined as greater than or equal to 30 consecutive amino acids). The human proteome assembly was downloaded from UniProt (Proteome: UP000005640) in August 2019. Overrepresentation statistics were computed using the standard Fisher’s test with Bonferonni correction for multiple testing. The correction was applied in two steps to take into the account testing of multiple GO terms and testing of multiple clusters. The overrepresentation statistics and corrected p-values are available in **Supplementary Table 2**. In **Figure 3** and **Supplementary** Figure 5, the clusters are labelled with overrepresented GO term descriptions, which have been abbreviated and represented in a shorter form where applicable.

### Definition of functional categories and grouping of GO terms

To systematize the GO terms into broader categories, we consulted GO hierarchy (ancestor charts), GO subsets, and GO co-occurring terms. For GO overrepresented terms from the clustering analysis, we defined 23 broader functional categories (**Supplementary Table 2**). We applied the same strategy to group GO terms predicted to individual IDRs by FAIDR (see *Cross validation of FAIDR and functional annotations on the proteome level*). From 148 GO terms, we converged on 19 broader categories that we defined and enumerated as given in **Supplementary Table 6** and **Figure 4C**. For instance, we grouped subcomponents of a cellular location (e.g., nuclear lumen, nuclear pore complex, nucleolus, nucleoplasm) into a higher order category “Nucleus, #7”. While such decisions were relatively straightforward for cellular locations, some functions and processes were more challenging to neatly subscribe to only one broad category. For instance, we grouped “GTPase regulator activity” term with other cellular signaling related terms into category “Signaling, #8”, based on its frequent co-occurrence with the signaling related terms. However, when looking at the ancestor chart for the term, it is apparent that it could be alternatively joint with terms related to “enzyme activity” or “catalysis” into a different broader category. This example illustrates that our categorization features an inevitable degree of subjectivity and will be subject to future updates and improvements.

### Validation of FAIDR performance on independent datasets of disease-associated genes

To validate the performance of FAIDR on autism-spectrum disorder (ASD) associated genes, we utilized a leave-one-out approach. We iteratively trained the model on IDRs from 101 out of 102 ASD risk genes defined by (Satterstrom et al. 2020) (hereafter ‘ASD risk dataset’). We under sampled the negative data set to include IDRs from 400 randomly selected proteins from the human proteome. Based on the ROC from testing on held-out data, we defined the threshold value of likelihood that was used to assess predictions of ASD-risk proteome-wide (**Supplementary Table 7**). The top 10% of predictions were cross-validated against the SFARI (Simons Foundation Autism Research Initiative) dataset (Abrahams et al. 2013). This allowed us to evaluate the predictive power of FAIDR on an independent dataset specifically focused on ASD-associated genes.

Additionally, to assess the performance of FAIDR on genes associated with cancer, we also used two separate datasets from the COSMIC database: Cancer census and Cancer classic (Sondka et al. 2018). Firstly, we curated the positive class of Cancer census dataset by eliminating any redundant instances of IDRs that were also present in the positive class of the Cancer classic dataset. Subsequently, we utilized the Cancer classic dataset with 409 IDRs in the positive dataset to train FAIDR and the Cancer census dataset with 723 IDRs for testing.

### Prediction of condensate localization and feature specificity

To evaluate FAIDR’s effectiveness in predicting proteins associated with biomolecular condensates, we utilized two distinct datasets for each condensate category. The training datasets comprised 229, 165, and 519 proteins that were experimentally validated to localize to stress granules, nuclear speckles, and the nucleolus, respectively (Youn et al. 2019, Lu et al. 2019). The test datasets consisted of 32 (cytoplasmic stress granule, GO:0010494), 305 (nuclear speck, GO:0016607), and 234 (nucleolus, GO:0005730) unique proteins annotated for localization to the respective condensates in Gene Ontology Browser (https://www.ebi.ac.uk/QuickGO/). Proteins present in both training and test sets were excluded from the latter and retained in the former. Subsequently, FAIDR was trained on a total dataset of 2,917 proteins for stress granules, 2,224 for nuclear speckles, and 6,454 for the nucleolus. Finally, we assessed the performance of the model proteome-wide, which encompassed a total of 16,115, 16,778 and 12,578 for stress granule, nuclear speckles and the nucleolus, respectively.

### Code and data availability

This study made use of UniProt, ENSEMBL, PANTHER, SFARI, RNA Granule Database and Human Protein Atlas databases, as specifically referenced throughout. Code and example files to compute all the steps described in the methods are available on GitHub (https://github.com/IPritisanac/IDR_ES/). The hierarchically clustered evolutionary Z-scores of human IDRs (i.e., the functional map), tutorial on the exploratory and automatic analysis of the map, IDR clusters, IDR-ome sequence and alignment files, FAIDR t-statistic and target files for top predicted GO terms are available at Zenodo (https://zenodo.org/records/10812875).

## Acknowledgements

IP and TRA were supported by a LiUNA! Fellowship for Research Innovation from The Hospital for Sick Children and a Banting Postdoctoral Fellowship from the Canadian Institutes of Health Research (CIHR), respectively. AMM and JDF-K acknowledge support from the CIHR (CIHR Foundation Grant (grant no. FDN-148375) to JDF-K; CIHR grant no. PJT-148532 to AMM and JDF-K) and the Canada Foundation for Innovation (CFI) for funding to AMM. JDF-K holds a Canada Research Chair in Intrinsically Disordered Proteins. AMM holds a Canada Research Chair in Computational Biology. We thank Ozren Kisić Morduš for creating the artwork in Figure 6.

