## Supplementary Information for "A Functional Map of the Human Intrinsically Disordered Proteome"

### Supplementary Tables

**Table S1**: Additional features used to compute human evolutionary signatures, in addition to those defined by Zarin et al. eLife, 2019)

**Table S2**: Overrepresented GO terms and molecular features within clusters extracted either through exploratory analysis (Tabs A-C) or automatic cluster analysis (Tab D) from the comprehensive map of the human IDR-ome. Clusters are named based on enumeration.

**Table S3**: IDRs in clusters that were extracted from the global map of the human IDR-ome (related to Figure 3). The IDR identifier comprises the UniProtID of the protein, followed by the first and last amino acid of the IDR, separated by the symbol "_". Tabs A-F), IDRs from selected clusters, highlighted in the Figure 3B and discussed in the text. Tab (G) IDRs from all clusters extracted at a distance threshold.

**Table S4**: IDPs in the human proteome and in the map of the IDR-ome. The UniProt IDs for proteins that are 95% or more disordered (predicted by SPOT-Disorder v1) are listed in the first tab. Of these IDPs, those that are clustered in the map of the IDR-ome (related to Figure 3) are listed in the second tab, and the final tab lists IDPs that are in clusters with overrepresented GO terms.

**Table S5**: Proteins of unknown function in the map of the IDR-ome. The first tab lists UniProt IDs for proteins of unknown function, as defined by the neXtProt Functional Proteome Project. The second tab lists the UniProt IDs for the proteins that contain IDRs, as predicted by SPOT-Disorder v1. The third tab lists the UniProt IDs for IDR-containing proteins of unknown function that are clustered in the map of the IDR-ome (related to Figure 3), and the final tab lists those that are in clusters with overrepresented GO terms.

**Table S6**: The results of the testing of FAIDR models, which predict the association of human IDRs with various Gene Ontology (GO) terms, as related to Figure 4. During training, the FAIDR models utilized GO term annotations at the per-protein level. Specifically, a distinct FAIDR model was trained for each of the 601 GO terms enumerated in Tab A). To evaluate classification performance, bootstrap test samples were employed, and assessments were made at the per-protein level.

**Table S7**: Disease-risk and biomolecular condensates. Tabs A-D) Clusters from the global map of the human IDR-ome featuring overrepresentation of disease-risk genes or proteins involved in formation of biomolecular condensates. Tabs E, F) FAIDR predictions for disease-risk genes and proteins in biomolecular condensates based on evolutionary Z-scores of IDRs.

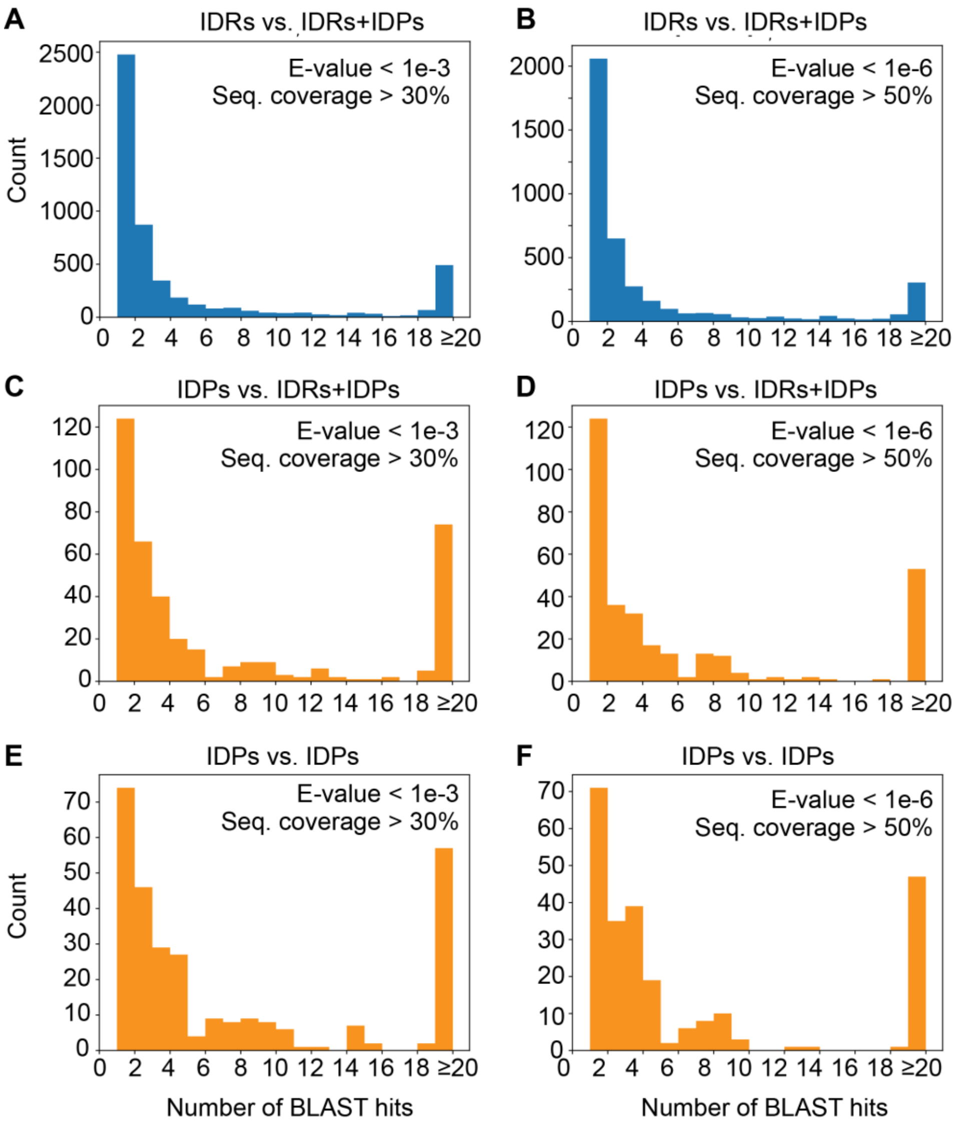


**Supplementary Figure 1.** Sequence similarity of the human IDR-ome as detected by BLAST. (**A**, **B**) Each IDR sequence in the IDR-ome was subjected to a BLAST query using the entire IDR-ome (including IDPs) as the sequence search database. The histograms show the number of BLAST hits that were detected given the threshold E-value and query sequence coverage of < 1e-3 and > 30% (**A**) or < 1e-6 and > 50% (**B**), shown in the upper right corner. In (B), only 21% of IDR sequences yield a BLAST hit, and 80% of the IDRs with hits have fewer than 5 hits. (**C, D**) When only fully disordered proteins (IDPs) are considered using the same IDR-ome search library, a higher proportion of IDP sequences return BLAST hits (44%) yet 91% of these IDPs have fewer than 5 hits. (**E, F**) A similar situation is observed when IDP sequences are subjected to a BLAST search using only IDPs as the sequence search library (34% have BLAST hits, of which 68% of these have five or fewer hits). However, fewer BLAST hits are returned overall as compared to panels C and D, suggesting that some IDPs and IDRs are similar.

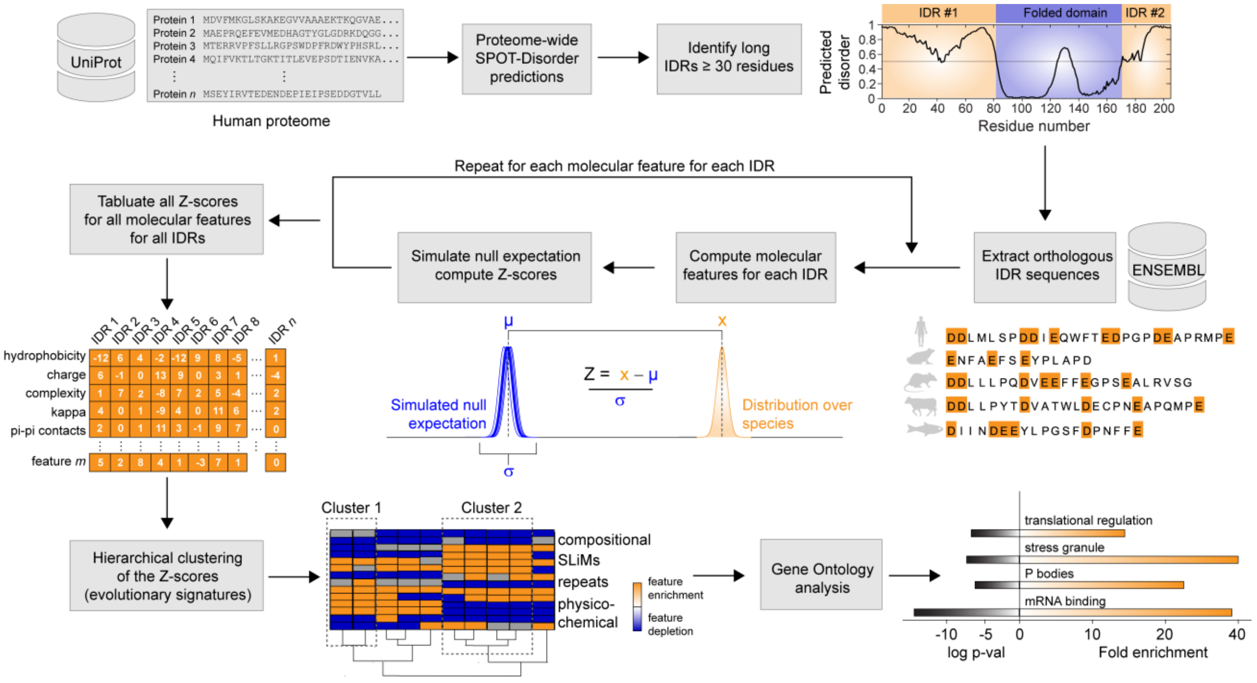


**Supplementary Figure 2.** Flow chart outlining the methodology used to compute evolutionary conserved molecular features of human IDRs. Protein sequences from the human proteome were obtained from UniProt and IDRs were identified with SPOT-Disorder. Only the IDRs longer than or equal to 30 consecutive residues were considered. Orthologous IDR sequences were extracted from the ENSEMBL database. A range of molecular features (Zarin et al. 2019, Table S1) were computed for each IDR and distribution of values of each feature over orthologous IDRs was compared to that of the simulated null expectation. The Z-scores were tabulated for each molecular feature for each human IDR. The set of all Z-scores of an IDR constitutes an evolutionary signature of the IDR, a term previously proposed by Zarin et al. for yeast IDRs. The evolutionary signatures are hierarchically clustered and the resulting tree structure subsequently segmented to branches that extract groups of IDRs with similar evolutionary signatures to their respective clusters. Each cluster is subjected to a Gene Ontology and feature enrichment analysis (see text for more details).

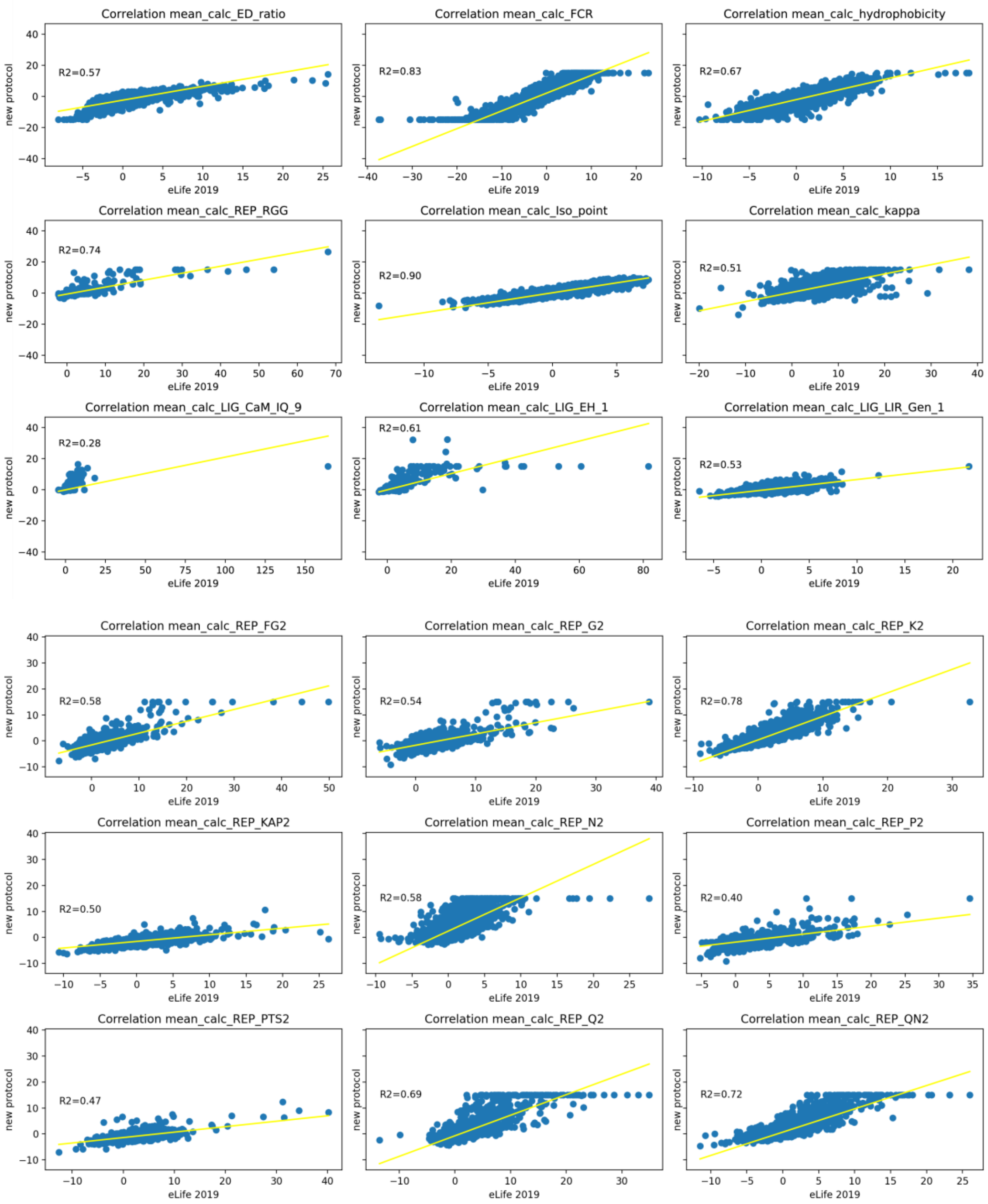


**Supplementary Figure 3.** Comparison of evolutionary Z-scores of yeast IDRs (*S. cerevisiae*) published by Zarin et al. (Zarin *et al.* 2019, x-axis) to those recomputed with the current protocol for evolutionary signatures (y-axis), which was used in this study to derive the human evolutionary signatures. The square of the Pearson correlation coefficient ($R^2$-value) from the linear least-squares regression is displayed for each comparison. The two protocols show a good agreement for most features. Note that the evolutionary signatures computed with the current protocol are truncated at values +/-15.

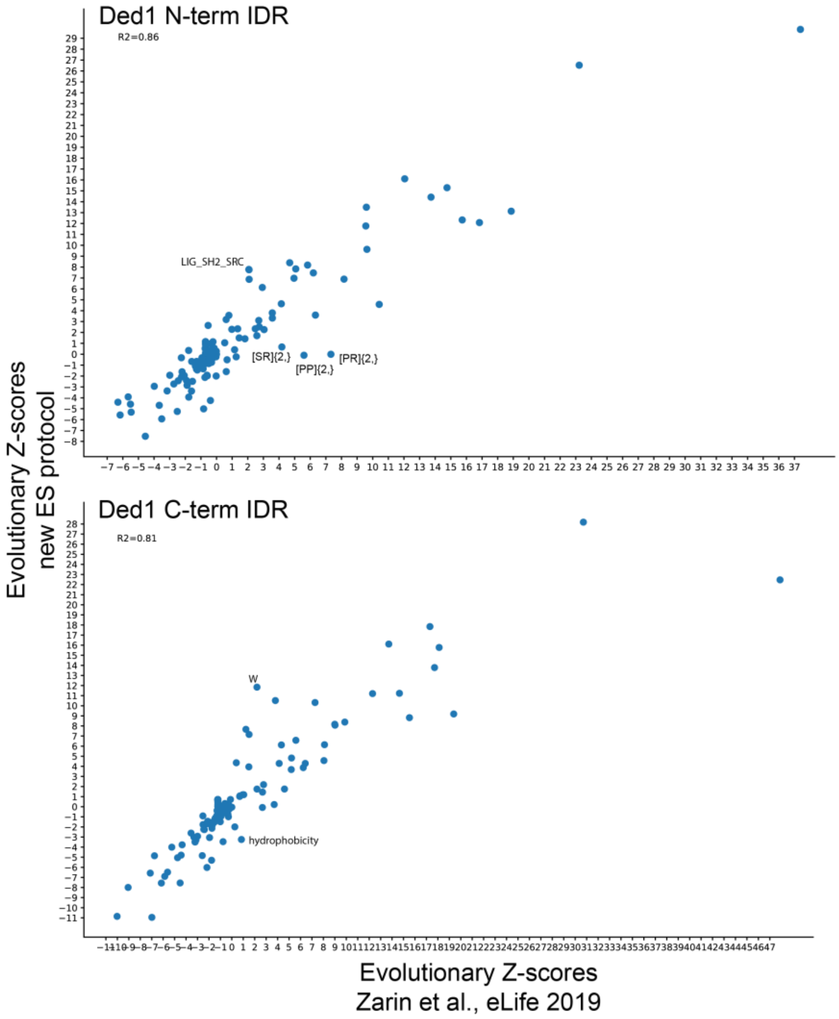


**Supplementary Figure 4**. Comparison of evolutionary Z-scores of yeast Ded-1 N- (top) and C-terminal (bottom) IDRs computed by Zarin *et al.* (eLife 2019, x-axis) to those recomputed with a simplified (new) protocol for evolutionary signatures (y-axis). The square of the Person correlation coefficient ($R^2$-value) is displayed for both comparisons ($R^2$ = 0.86, 0.81). A good agreement between the two protocols is found for most features. A few features that show larger deviations in the computed Z-scores between the two protocols are indicated on the plots. The deviations were selected for display when the Z-score of a feature would not be considered significant by one of the protocols (|Z-score|<3), but would be found significant by another. Small deviations in exact Z-scores of features for which both protocols agree on significance is not marked.

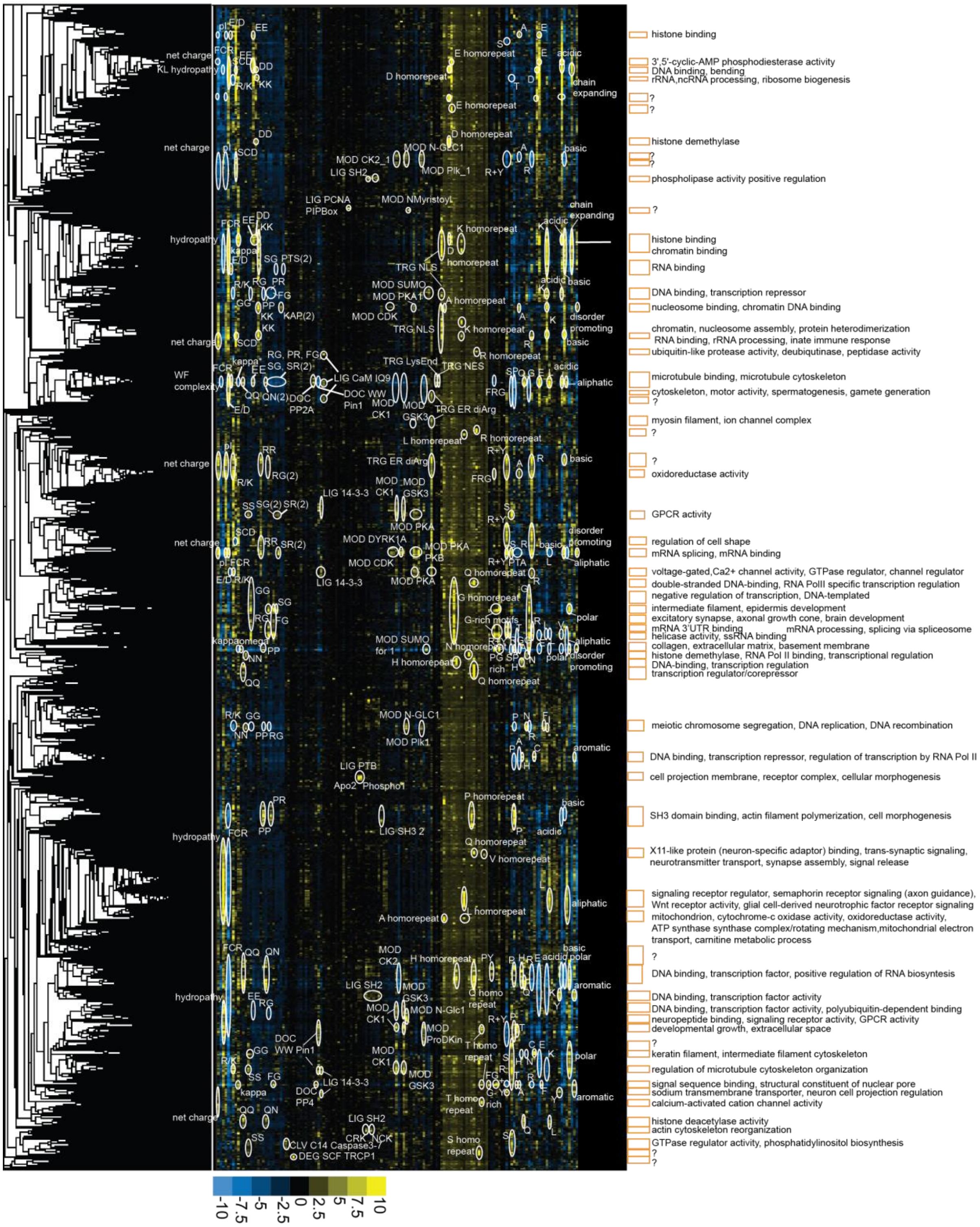


**Supplementary Figure 5.** A global map of human IDRs obtained through hierarchical clustering of evolutionary conserved molecular features. Hierarchical clustering of the human IDR-ome using evolutionary signatures with features defining various clusters annotated with white circles. A positive or negative Z-score, respectively, is defined by a higher or lower value of a mean of feature over orthologous IDRs than would be expected in the absence of evolutionary conservation. Orange rectangles on the right indicate positions of selected clusters with molecular functions, biological processes and/or sub-cellular localizations that have statistically significant overrepresentation in the cluster listed. Question marks denote the clusters strongly defined by evolutionary Z-scores, but with no known functional annotations.

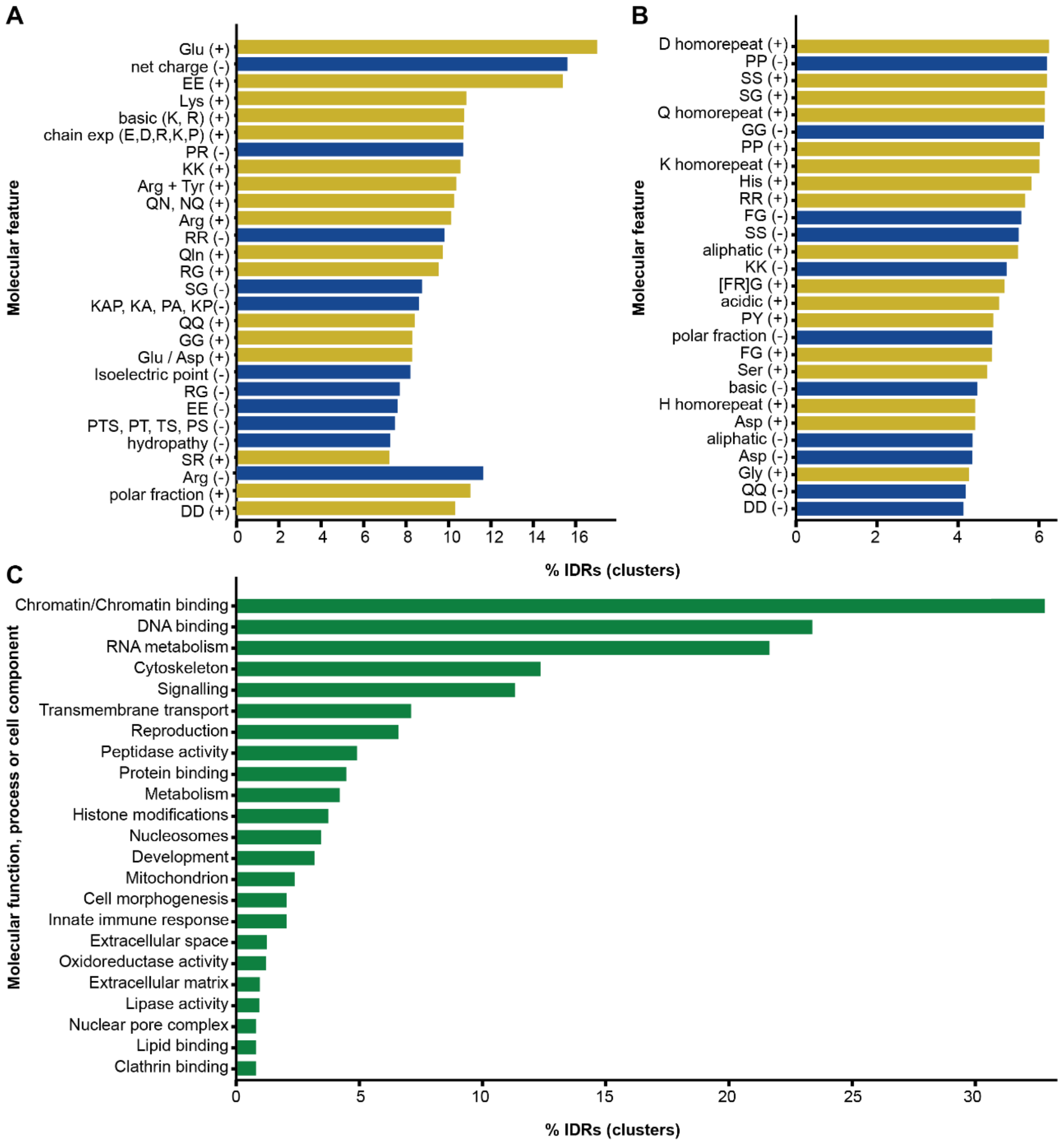


**Supplementary Figure 6.** Dominant molecular features and biological functions in the global map of human IDRs. **(A, B)** Molecular features that are significantly overrepresented by positive (yellow) or negative (blue) Z-scores in 93 clusters extracted from the global map through exploratory analysis. The clusters and their defining features are highlighted throughout the map shown in **Supplementary Figure 5** with orange squares and white circles, respectively. A subset of clusters that featured overrepresentation of specific GO terms (53/93) is listed in **Supplementary Table 2 (Tab A)**. **(C)** The GO-term overrepresentation from IDRs in the 53 clusters, as depicted in **Supplementary Figure 5** and documented in **Supplementary Table 2** (**Tab A**), were assigned to broad functional terms as given in **Supplementary Table 2 (Tab B)**. In all panels, the x-axis reflects the percentage of IDRs that are allocated to the clusters, and not the total percentage of IDRs in the proteome.

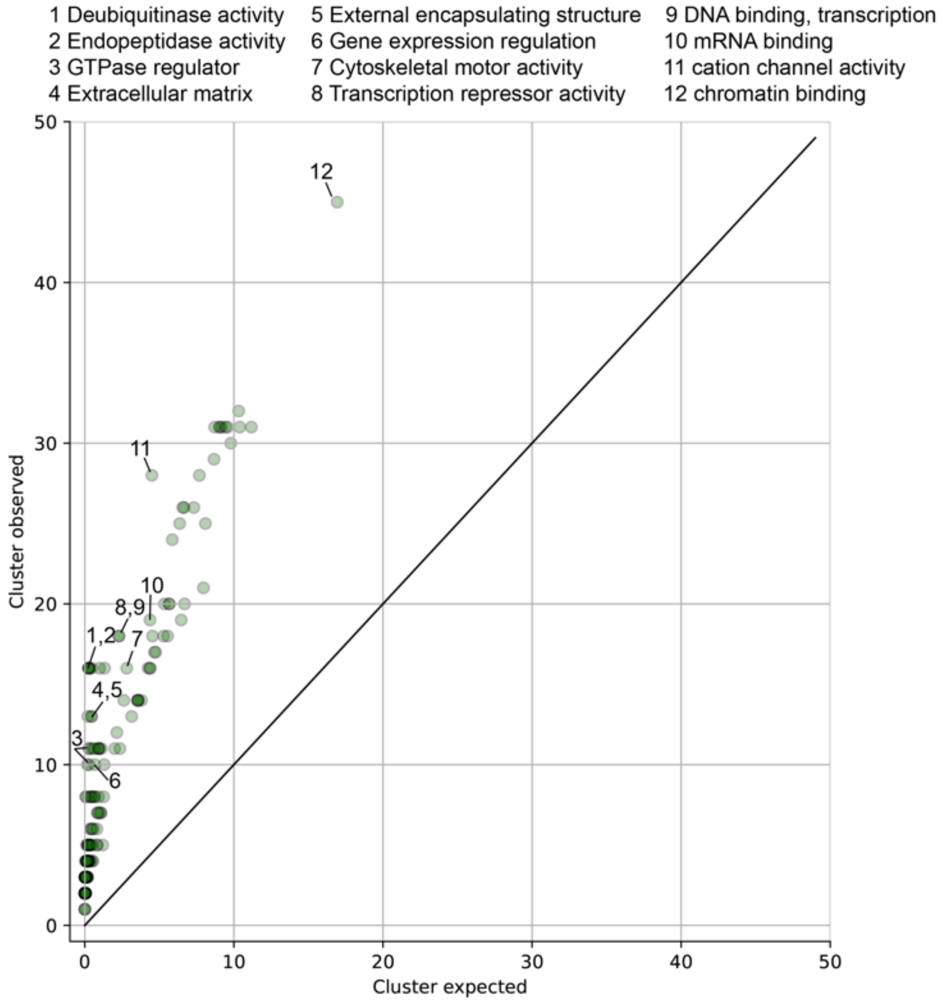


**Supplementary Figure 7.** Overrepresentation of GO-terms in 393 IDR clusters extracted from the global map through automated analysis using a clustering threshold of 0.7. This threshold corresponds to the uncentered correlation distance between vectors representing evolutionary Z-scores of human IDRs. Some of the GO terms most deviating from expectation have been enumerated 1-12. Complete data are available in **Supplementary Table 2** (**Tabs D&E**).

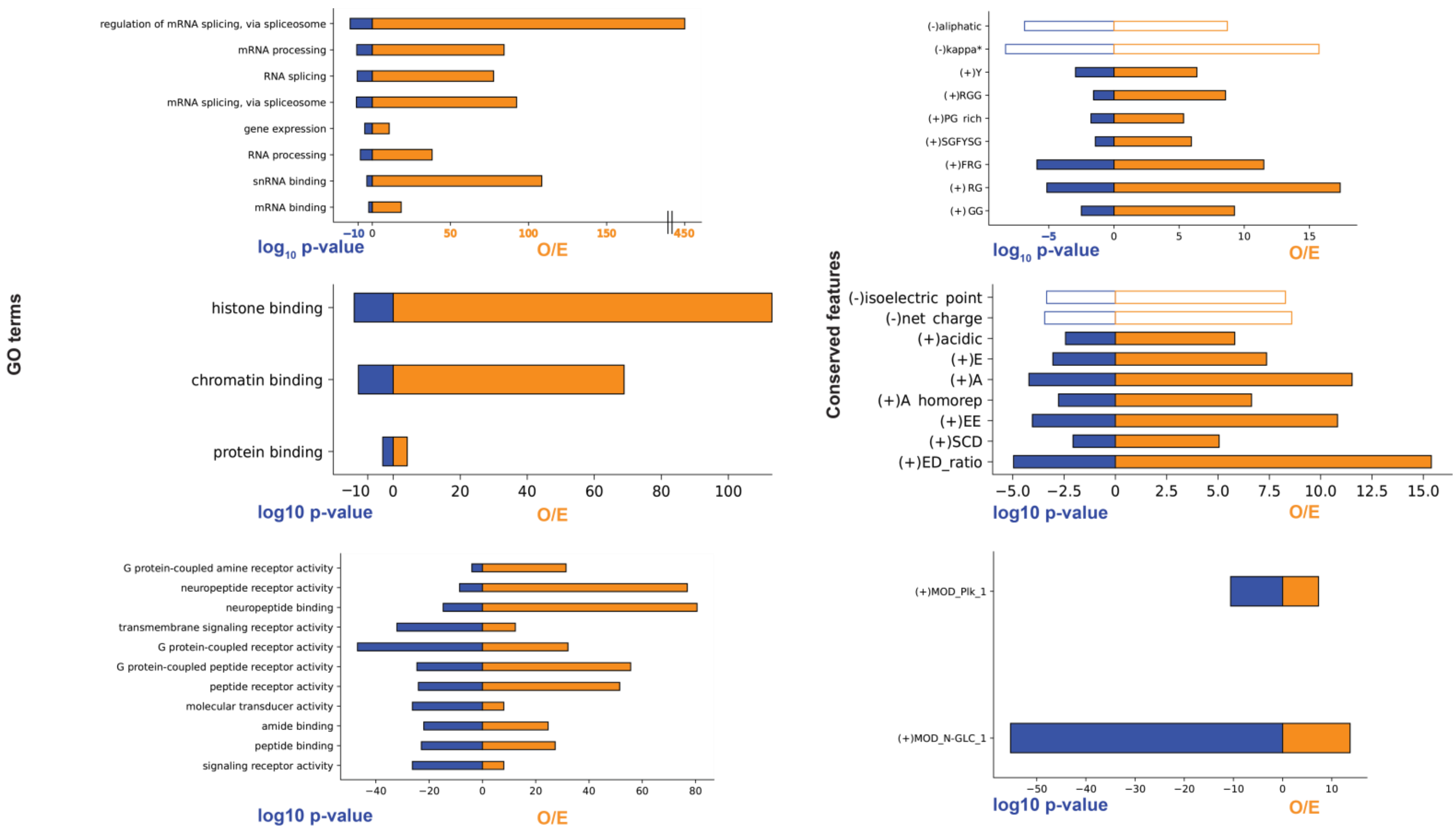


**Supplementary Figure 8.** A summary of GO term overrepresention and its statistical significance for a selection of clusters from the global map of the human IDR evolutionary signatures (see also **Figure 3** and **Supplementary Table S2**). The overrepresentation is calculated as a ratio of the observed versus expected number of proteins (genes) annotated to the function under the GO-term in the select cluster. The p-values are calculated based on a Fisher's exact test and corrected for multiple comparisons using a Bonferonni correction method. In the conserved features plot (right side), open and filled bars correspond to features with positive and negative Z-scores, respectively.

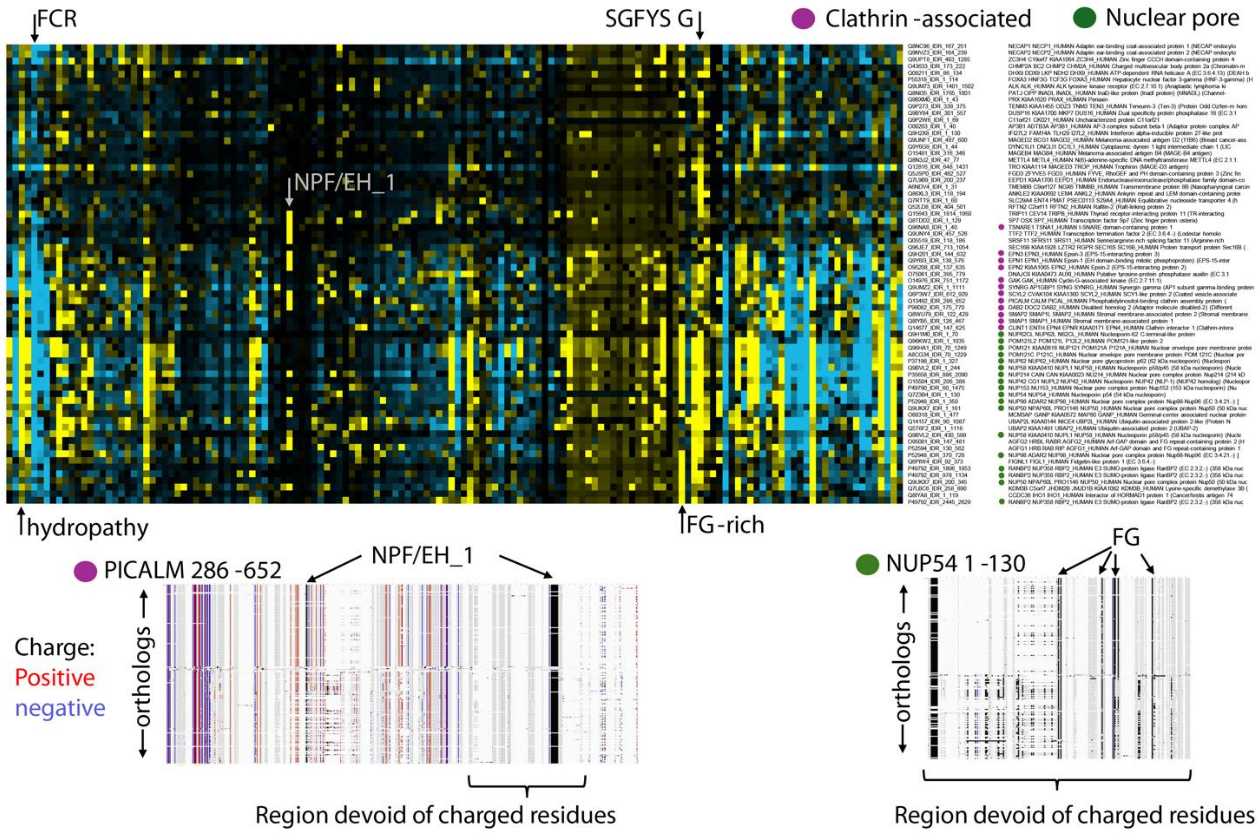


**Supplementary Figure 9.** Comparison of the evolutionary signatures for IDRs that are associated with nuclear pores and clathrin-associated endocytosis. Evolutionary signatures for 69 IDRs are shown as rows. Important molecular features for the clustering of these IDRs are indicated, such as the fraction of charged residues (FCR), hydropathy, Gly content (G), Phe-Gly (FG) motifs, and Ser/Gly/Phe/Tyr/Ser (SGFYS) motifs. The IDRs, UniProt IDs, and protein names are listed on the right side. Green circles indicate proteins that are associated with the nuclear pore, while purple circles denote those associated with clathrin-mediated endocytosis. In the bottom panel, sequence alignments are shown from a representative clathrin-associated IDR (PICALM residues 286-652) and nuclear pore-associated IDR (NUP54 residues 1-130). The first row is the amino-acid sequence of the respective human IDR, and all following rows correspond to orthologous sequences. Gaps are indicated in white while unchanged amino acids appear as grey columns and positive and negatively changed amino acids are colored red and blue, respectively. In the PICALM 286-652 alignment, the NFP/EH_1 motif matches are highlighted with black (indicated with arrows) and in the NUP54 1-130 alignment FG motif matches are highlighted with black . Regions that are devoid of charged residues are also indicated. See the main text for more details regarding the significance of these motifs and molecular features.

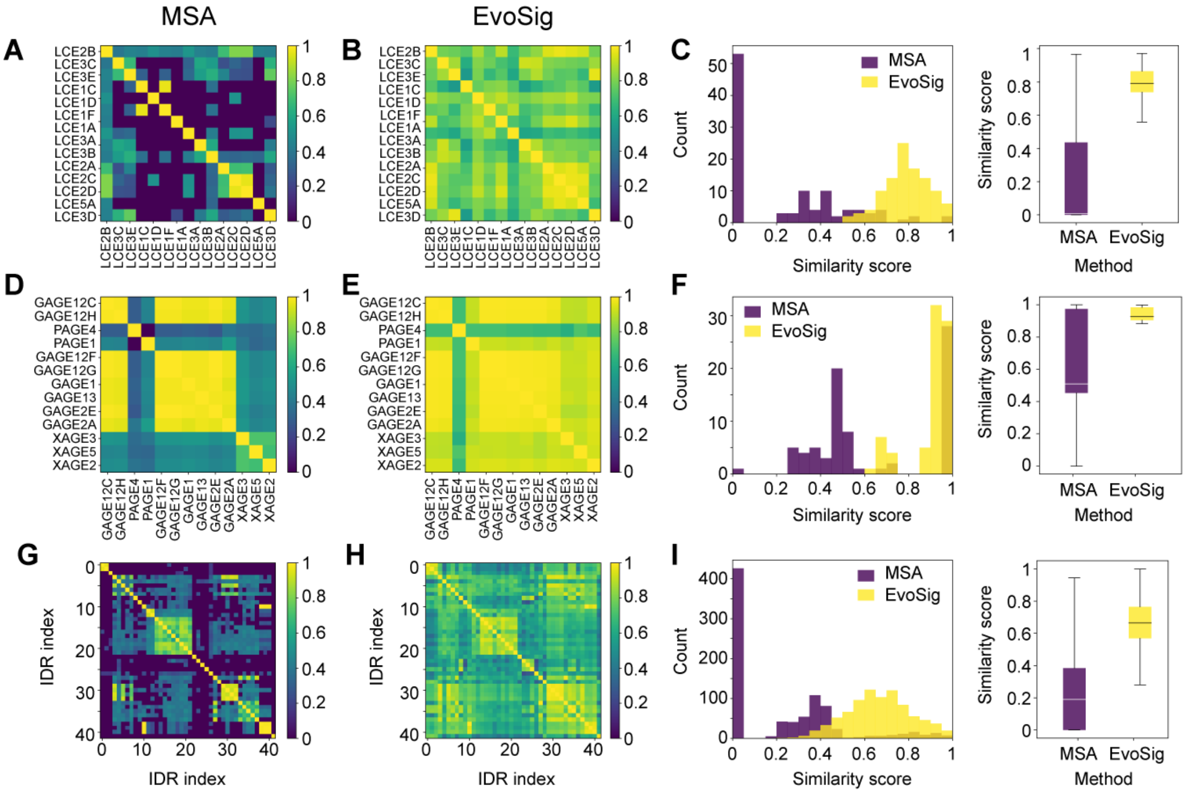


**Supplementary Figure 10.** Clusters enriched in IDP families cannot be explained by positional alignment alone. IDPs from three different clusters from our map of the human IDR-ome are shown in panels (**A, B, C**), (**D, E, F**), and (**G, H, I**), respectively. Panels (**A, B, C**) depict the cluster with 17 IDRs of which 15 are IDPs corresponding to members of the late cornified envelope (LCE) family of proteins. Panels (**C, D, E**) show the cluster with 21 IDRs of which 13 are IDPs from the G-, P-, and X-antigen (GAGE, PAGE, XAGE) families of proteins. Panels (**F, G, I**) correspond to the cluster of 76 IDRs of which 42 are IDPs, with 35 of these being from the keratin-related (KR) family of proteins and the other 7 from the small glycine and cysteine repeat (SGCR) family of proteins. The axes of these panels simply depict numbers between 1 and 42 for clarity. The first column (**A, D, G**) depicts heat-map representations of sequence similarity between the IDPs as measured by positional conservation (MSA). The plotted values are the product of the alignment coverage (fraction of the query sequence length) and the sequence identity over the aligned region. If any sequence did not align to at least 50% of the query sequence with at least 30% sequence identity, a value of 0 was assumed. Off-diagonal values that are closer to 1 indicate more similar sequences. The second column (**B, E, H**) shows the cosine similarity metric (**Methods**) between vectors in the 144-dimensional evolutionary signature space (EvoSig). Values that are closer to 1 signify that two sequences are closely related as measured by evolutionary signatures. For all three clusters, the evolutionary signature approach shows that all IDPs within the cluster are related whereas the alignment approach yields many IDPs that appear unrelated to others. The third column (**C, F, I**) shows histograms displaying the values (Similarity scores) that were extracted from the upper triangle of the heat maps shown in the preceding two columns. The diagonal elements were excluded. The MSA and EvoSig approaches are colored purple and yellow, respectively. To the right of the histograms is a box plot showing the spread in similarity scores for the MSA and EvoSig approaches. The box extends to the upper and lower quartiles, and the line within the box corresponds to the median value. The lines outside the box correspond to data within 1.5-fold the interquartile range, and outliers beyond these values were excluded from the plot.

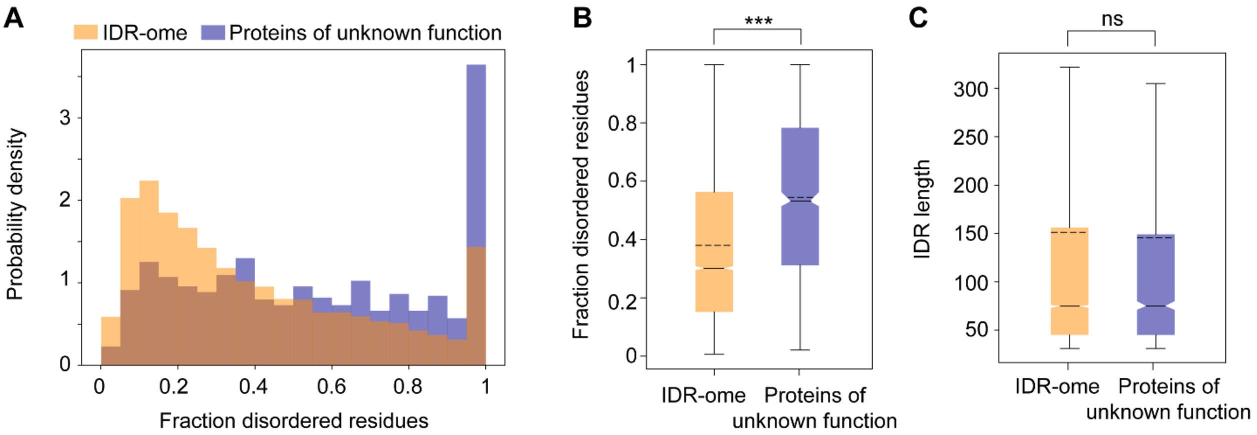


**Supplementary Figure 11.** Disorder content within human proteins of unknown function. (**A**) Histogram of SPOTD-predicted disorder content in proteins of unknown function (blue). The x-axis depicts the fraction of disordered residues (disordered residues / total residues) and the y-axis is the probability density. The corresponding histogram for proteins in the human IDR-ome is shown in orange. (**B**) Box plot representation of the data in panel A with the same color scheme. (**C**) Box plot representation of IDR lengths in the IDR-ome (orange) compared to the set of proteins of unknown function (blue). Distributions were compared with the Mann-Whitney test for statistical significance. Asterisks (***) indicate p-value < 0.001. . The box extends to the upper and lower quartiles, and the line within the box corresponds to the median value. The lines outside the box correspond to data within 1.5-fold the interquartile range, and outliers beyond these values were excluded from the plot.

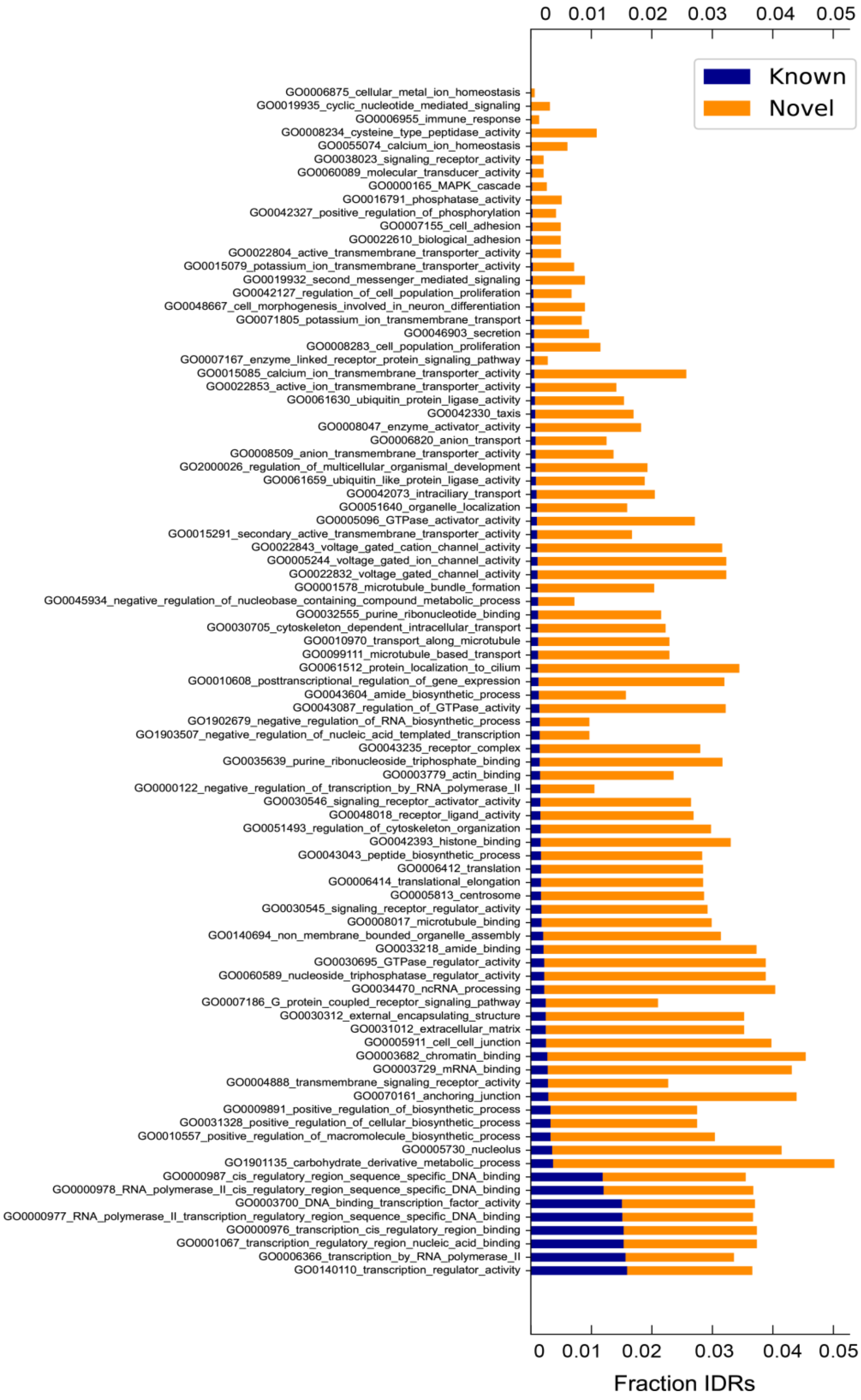


**Supplementary Figure 12.** The mapping of functional categories, which are summarized in Figure 4, to individual GO terms (Y-axis), upon which FAIDR was trained and the subsequent functional annotation is based.
